# Mapping of individual sensory nerve axons from digits to spinal cord with the Transparent Embedding Solvent System

**DOI:** 10.1101/2021.11.13.467610

**Authors:** Yating Yi, Yi Men, Shiwen Zhang, Yuhong Wang, Zexi Chen, Ed Lachika, Hanchuan Peng, Woo-Ping Ge, Hu Zhao

**Affiliations:** Chinese Institute for Brain Research, Beijing, P.R. China; Sichuan University, College of Stomatology, Chengdu, Sichuan, P.R. China; Intelligent Imaging Innovations (3i) Inc., 3509 Ringsby Court, Denver, CO 80216, USA; SEU-ALLEN Joint Center, Institute for Brain and Intelligence, Southeast University, Nanjing, 210096, P.R. China

**Author notes:** These authors contributed equally (Yating Yi, Yi Men, Shiwen Zhang, Yuhong Wang).

**Keywords:** tissue clearing, connectome mapping, peripheral nervous system, dorsal root ganglion, transparent embedding, TESOS

## Abstract

Understanding the connections and projections of neurons has been a fundamental issue for neuroscience. Although strategies have been developed to map the projection of individual axons within the mouse brain, high resolution mapping of individual peripheral nerve axons in peripheral organs or spinal cord has never been achieved. Here, we designed the Transparent Embedding Solvent System (TESOS) method and developed a technical pipeline for imaging, reconstructing and analyzing large samples containing various tissue types at sub-micron resolution. The mouse whole body was reconstructed at micron-scale resolution. We were able to image, reconstruct and analyze the complete axonal projection of individual sensory neurons within an intact mouse paw or spinal cord at sub-micron resolution. Furtherly, we imaged and reconstructed the entire projection course of individual sensory neurons from adult mouse digits to the spinal cord. The TESOS method provides an efficient tool for micron-scale connectome mapping of the peripheral nervous system.

## Introduction

Mammalian neurons give rise to extensive axons which may travel a very long distance to convey information across different areas. Mapping neural connectivity at single axon level is crucial for understanding the neuron properties and the routing of information flow. Sensory neurons are the central components of somatosensory system. Somas of sensory neurons reside in the dorsal root ganglion (DRG) and trigeminal ganglia. They extend one axonal branch to the skin and connect with sensory endings to perceive external stimuli (Zimmerman et al., 2014) and another axonal branch to the spinal cord or the brainstem to convey the information to the central nervous system (Abraira and Ginty, 2013).

Most neural connectome research has been focused on brain. Various strategies have been designed to image and map axon connectivity in brain at single axon level (Economo et al., 2016; Gong et al., 2016; Li et al., 2010; Zeng, 2018; Zheng et al., 2013). Connection and projection mapping of peripheral nervous system were much less studied. Connectivity mapping of single sensory neurons in peripheral organs or spinal cord at high resolution has never been achieved in rodent model (Kuehn et al., 2019). Unlike brain, sensory axons travel a very long distance across complex tissue types including skin, muscle, fat and even bone etc., which made the high-resolution imaging or tracing very difficult. Thick histological sections or flat-mounted sample remained to be the only available approaches for investigating the peripheral or central projections of sensory neurons. which provided limited spatial information(Browne et al., 2020; Kuehn et al., 2019; Olson et al., 2016; Woodbury et al., 2001).

Tissue clearing has been a major technical breakthrough for microscopic imaging. By treating biological samples with various chemicals, tissue could be turned into transparent. Current clearing methods can be largely divided into three major categories including aqueous methods, solvent based methods and hydrogel-based methods (Tainaka et al., 2016). Most tissue clearing methods followed similar chemical principles and were comprised of steps including fixation, decalcification (for hard tissue), decolorization, delipidation, dehydration (for solvent based clearing methods only) and RI matching (Tainaka et al., 2016). Tissue clearing technique provides a powerful approach for optical imaging deep within biological specimens. Tissue clearing have also been employed to investigate peripheral nerves travelling long range within mouse body (Cai et al., 2019). In combination with nanobody staining and light sheet microscope, single axon resolution was achieved in some regions including the skin and the muscles surfaces (Cai et al., 2019).

To achieve isotropic high resolution throughout the entire sample is the major challenge for all current tissue-clearing based imaging. Even in cleared organs, when the optical path is long, aberration still builds up due to accumulated refractive index (RI) mismatch alone the long optical path and leads to resolution deterioration (Ueda et al., 2020). Optical aberration in peripheral organs and tissue is even mor severe than in the brain due to varied tissue components. Another technical barrier is the microscope objective. Image resolution is physically determined by the numeric aperture (NA) number of the objective. A high NA objective always has a very short working distance, which limited its image depth within the tissue (Chen et al., 2020; Gao, 2015).

In the current study, we designed the Transparent Embedding Solvent System (TESOS) clearing method and built a technical pipeline to image, reconstruct and analyze sensory neuron axons projections at sub-micron resolution. UV light-initiated polymerization reactions transform cleared tissue together with the clearing solution into transparent organogel, a process we named *transparent embedding*. Samples remained fully transparent with unchanged endogenous fluorescence and the mechanical strength was enhanced by >150 folds. We designed microtome or milling motor-based processing platform and imaged samples with a block face imaging strategy. We were able to achieve sub-micron resolution in large samples composed of various tissue types. With the TESOS method, we reconstructed and analyzed peripheral and central projections of single sensory neuron axons within adult mouse forepaw or spinal cord. We also reconstructed and analyzed the entire projection course of five sensory axons extending for ∼6cm from adult mouse forepaw to DRG and spinal cord. Analysis of our data provided valuable information on the central projection pattern of sensory neurons.

## Results

### Development of the Transparent Embedding Solvent System (TESOS)

The TESOS method is a solvent based tissue clearing method designed based on our previous PEGASOS method (Jing et al., 2018). It preserved the advantage of potent clearing capability for both hard and soft tissues and incorporated the concept of “*transparent embedding*”. The TESOS method is consisted of multiple steps including fixation, decalcification (for hard tissue), decolorization, delipidation, dehydration and clearing (**Figure 1 A**). After 4% PFA fixation, 25% *N,N,N*′,*N*′-Tetrakis(2-Hydroxypropyl)ethylenediamine (Quadrol) solution was used as the decolorizing reagent to remove heme (Jing et al., 2018; Tainaka et al., 2014). We applied gradient tert-butanol (tB) solution for delipidation. We designed tB-Quadrol reagent for dehydration, which is composed of 70% tB + 30% Quadrol. For final tissue clearing, we designed BB-BED468 clearing medium (RI 1.55), which is composed of 47%(v/v) benzyl benzoate (BB), 48% (v/v) of bisphenol-A ethoxylate diacrylate M_n_ 468 (BED468), 5% (v/v) Quadrol and then supplemented with 2% w/v of 2-Hydroxy-4’-(2-hydroxyethoxy)-2-methylpropiophenone as the UV initiator (**Figure 1A**). For organs containing hard tissue, decalcification treatment with EDTA solution was included before decolorization step (**Figure S1 A**). Typically, it took 8 (soft tissue organs) or 12 days (hard tissue organs) to reach final transparency. The clearing treatment rendered organs of various tissues to high transparency, including brain, mandible, mouse paw with skin, vertebrae with bone and muscles, spleen, liver, heart, femur or even whole body of mouse pups (**Figure 1D, Figure S1 B**).

**Figure 1.**
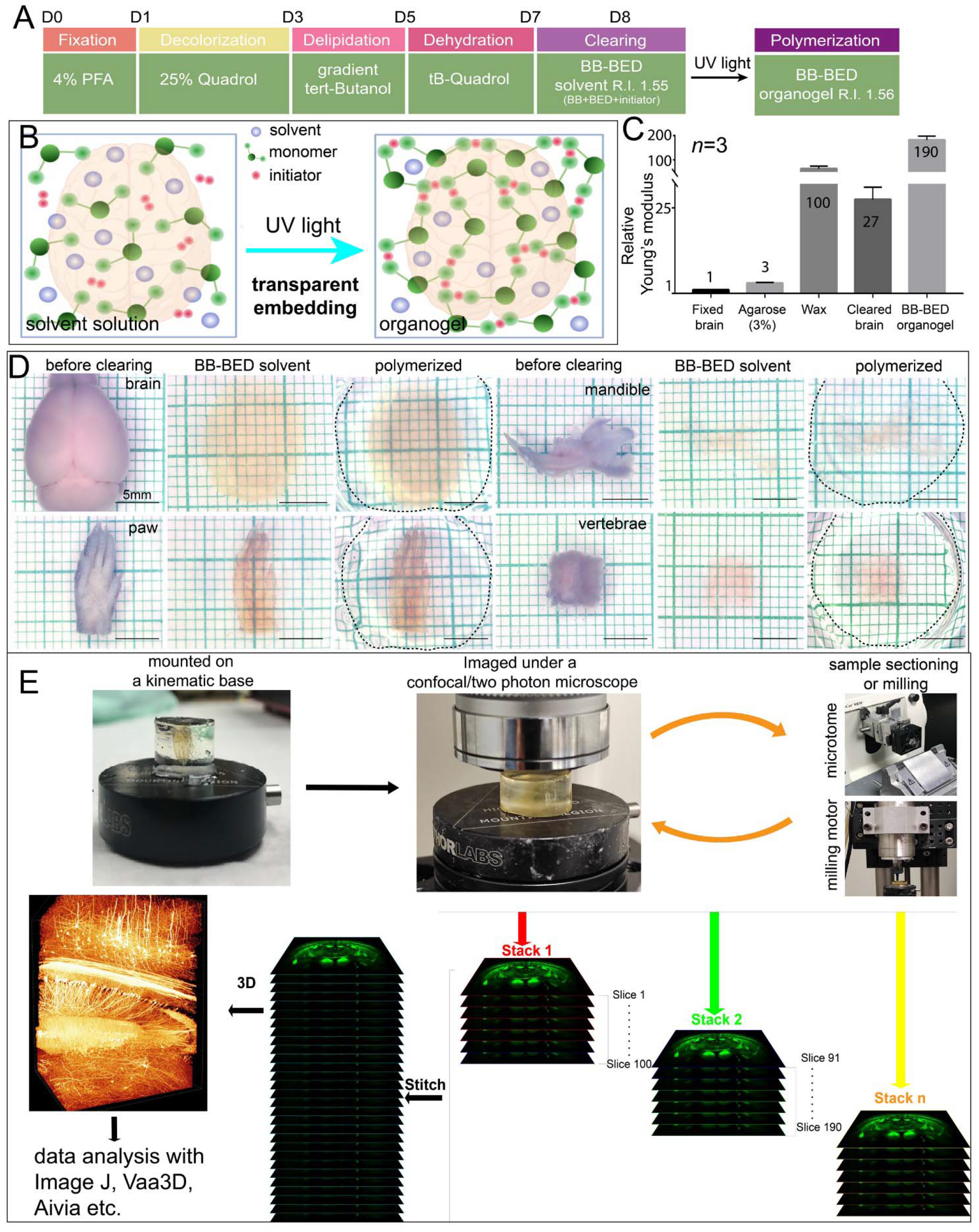
Technical pipeline of the TESOS method. (A). Treatment steps of the TESOS method for soft tissue organs. (B). Formation of the organogel is the conceptual basis for the process of transparent embedding. BB-BED clearing medium is mainly composed of solvent, monomer and UV initiator. UV light initiates polymerization and converts the solvent solution into organogel. (C). Relative mechanical strength of various samples displayed as the relative Young’s module. The value of freshly fixed brain was normalized as 1. (D). Images of brain, mandible bone, paw and vertebrae samples of adult mouse before clearing, after clearing and after polymerization. Dotted lines outline the boundary of the organogel. (E). Imaging and data processing pipeline of the TESOS method. See also Figures S1 and S2.

Bisphenol-A ethoxylate diacrylate (BED) is a commonly used monomer in dental resin due to its mechanical strength and low-toxicity. It contains acrylate groups at two ends. In the presence of UV initiator and UV light, BED molecules could be cross-linked through free radical polymerization reactions. Benzyl benzoate (BB) solvent did not participate in the polymerization reaction and was dispersed among the crosslinked polymer chains to form the organogel (**Fig 1 B**). The BB-BED organogel not only was transparent by itself, but also maintained the tissues within equally transparent as in the solution (**Figure 1 D, Figure S1 B, C)**.

To evaluate the endogenous fluorescence change, we performed quantified assays using brain samples harvested from *Thy1*-*EGFP* mice (P60). Samples were processed following either PEGASOS or TESOS method. Fluorescent intensity after PFA fixation was normalized as one. At the end of PEGASOS or TESOS clearing process, GFP fluorescence retained ∼70% of the original intensity with no significant difference between the two methods (**Figure S1 D**). UV initiated polymerization process has no significant impact to the GFP fluorescence (**Figure S1 D**).

To evaluate the impact of transparent embedding on the tissue transparency, *Thy1-EGFP* brain slices (**Figure S1 E, F**) or *Cdh5-Cre*^*ERT2*^*;Ai14* mice mandible (**Figure S1 G, H**) samples were imaged before and after polymerization. With a Leica 20X/0.95 immersion objective, images were acquired at the same locations of various depth. Polymerization process made no detectable impact to the image quality in both brain and hard tissue samples, indicating the tissue transparency was well maintained after polymerization.

The Young’s module of brain samples was measured before and after transparent embedding. The Young’s module of a cleared brain was ∼25 fold stronger than a freshly fixed brain, possibly due to the dehydration process. A transparently embedded brain was over 150 folds stronger than the freshly fixed brain. In comparison, agarose strength is ∼3 folds stronger than a brain. A paraffin block is ∼100 folds stronger than a brain (**Figure 1 C**).

Therefore, the TESOS method efficiently rendered various types of tissue highly transparent and the transparent embedding treatment improved tissue strength by more than 150 folds while preserving transparency and endogenous fluorescence.

### Process of transparently embedded samples with sectioning or milling platform

Next, based on the TESOS method, we designed and built sample processing platforms to image samples of any size at high resolution with a block-face imaging strategy (**Figure 1 E**). The magnetic kinematic base was originally used for re-positioning optical elements with high precision and high repeatability (**Figure S2 A**). Transparently embedded samples were glued onto the top plate of the kinematic base (**Figure S2 A**). A kinematic base bottom plate was secured under an upright confocal/two photon microscope (**Figure S2 B**). Samples were mounted under the microscope through the magnetic force. The top surface of the samples was imaged like a regular sample slide. The imaging depth is determined by the working distance of the objective.

Samples were transferred between the microscope and a rotary microtome (**Figure S2 C**). After each sectioning, new Z stack was acquired with 10% overlapping with the previous stack. Magnetic base is the key element to ensure the samples being returned to identical positions after sectioning. We also designed alignment step to ensure the sectioning plane is parallel to the imaging plane (see details in the method part). This strategy is applicable for brain and large body trunk.

Alternatively, top surface of samples could be removed with a high-speed bur on the milling motor. The milling motor was built next to the microscope stand (**Figure S2 D, E, F**). Samples were moved along a linear guide between the motor and the microscope. The linear guide is the key element to ensure the sample being repositioned precisely after milling. This strategy is applicable for both soft tissue and complex tissues organs.

To evaluate if sectioning or milling distorted sample surface, we tested with brain, vertebrae, femur and complete head of adult mouse samples, which included various tissue types including skin, hair, muscle, bone and nerve. Images were first acquired at 400µm depth for embedded brain, vertebrae or femur samples (**Figure S2 G, H, I, J, K, L, before sectioning**). Next, 390µm thickness of tissue was sectioned with a microtome. Same samples were re-imaged at the same locations, which were 10µm below the sectioning surface (**Figure S2 G, H, I, J, K, L after sectioning**). Overlaying of pre- and post-sectioning images indicated overall organization and very fine structures like neuron dendrites remain unchanged after sectioning (**FigureS2 G, H, I, J, K, L overlay**). Similar comparison was performed on adult mouse head sample processed with the milling motor setup. Overall structures and fine cellular structures also remained unchanged after milling treatment (**Figure 2 M, N**).

**Figure 2.**
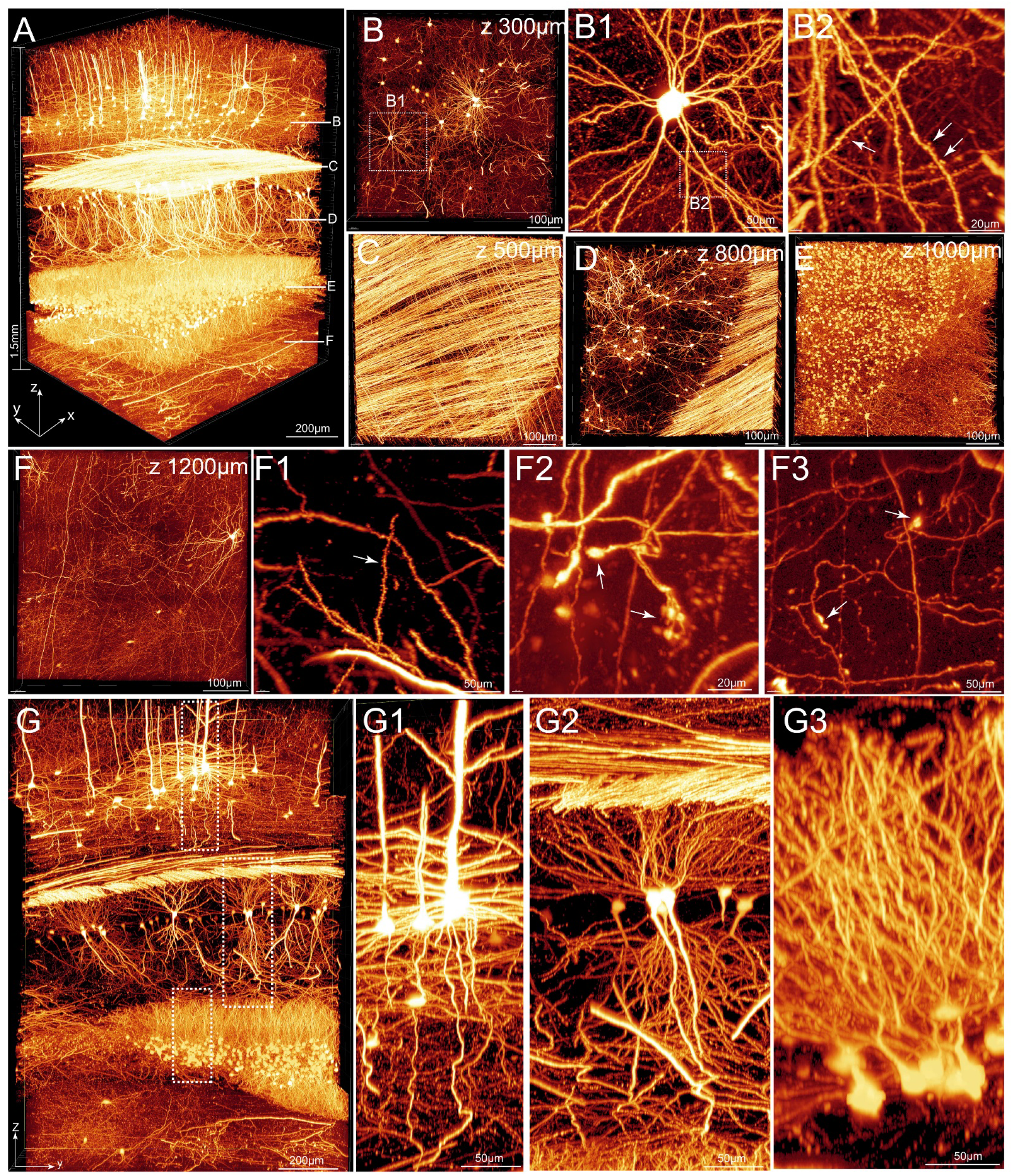
Sub-micron resolution imaging of *Thy1-EGFP* brain sample. Adult *Thy1-EGFP* mouse brain sample of 1mm (x) X 1mm (y) X 1.5mm (z) size was processed and imaged with a 40X 1.3NA objective (voxel size 0.26µm X 0.26µm X 1.2 µm). (A) The final image stack was stitched from 8 stacks with 240µm thickness for each stack. (B-F). Sub-blocks (size: x-1mm, y-1mm, z-0.2mm) were acquired at various Z positions indicated with white lines in (A) and displayed in x-y orientation. Boxed region in (B) was enlarged in (B1). Boxed region in (B1) was resliced and enlarged in B2 to display dendritic spines (arrows). Region in (F) was resliced and enlarged to display dendritic spines (arrow in F1) and boutons (arrows in F2 and F3). (G). A sub-block (x-200 µm, y-1mm, z-1.5mm) was displayed in y-z orientation. Boxed regions were enlarged to display descending neurons (G1), pyramidal neurons (G2) and granule cells (G3) respectively. See also Figures S3 and S4.

Sectioning or milling process did not cause detectable tissue distortion, which enabled adjacent image stacks to be stitched based on overlapping layers to reconstruct a complete image.

### Sub-micron resolution imaging of mouse brain, bone and vertebrae samples

We applied TESOS method on organs of different tissue types. *Thy1-EGFP* mouse brain sample of 1mmX1mmX1.5mm size was processed with the TESOS method and imaged with a Zeiss 40X/1.3NA objective with voxel size of 0.26 µm X 0.26 µm X 1.2 µm. The final image stack was stitched from 8 stacks (**Figure 2 A, Video 1**). Optical slices indicated high quality images were acquired at all z depth (**Figure 2 B-F**). Dendritic spines and boutons could be visualized at either 300µm or 1200µm depth (**Figure 3 B2, F1, F2, F3**). Y-Z optical slice indicated structure continuity in Z dimension was not disrupted by sectioning. Axons and dendrites remained continuous and intact in Z dimension (**Figure 2 G, G1, G2, G3**).

**Figure 3.**
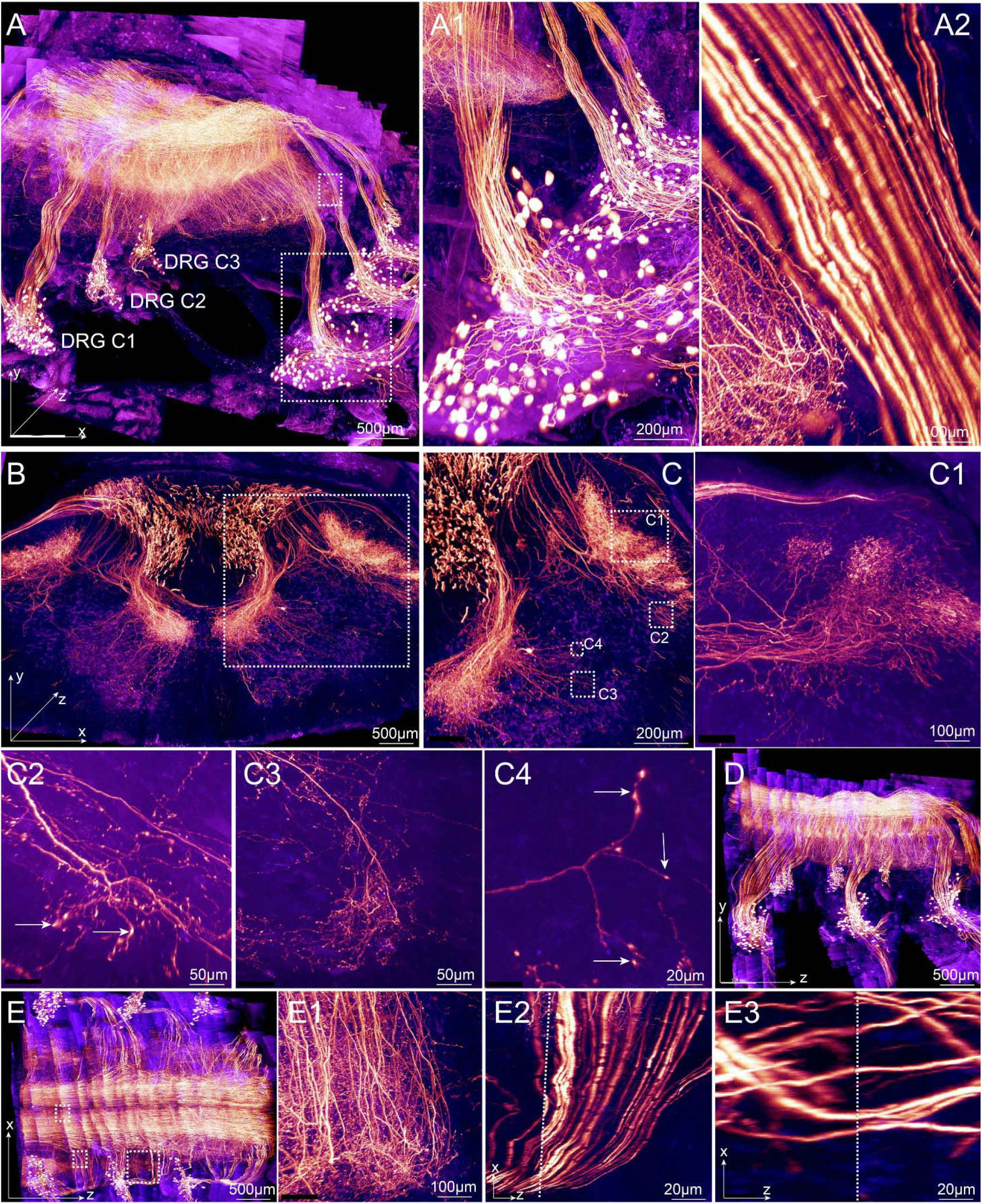
DRG sensory neurons and their projections in the spinal cord. Adult *Shh-Cre*^*ERT2*^*;Ai140* mouse vertebrae segment containing spinal cord, bones, attached muscle and skin (3.5mm X 2.2mm X 3mm) was processed for transparent embedding. A 40X/1.3NA oil immersion objective was used for imaging (voxel size 0.37µm X 0.37µm X 1.2 µm) (gold, GFP signal; blue, autofluorescence). (A). Cervical DRG pairs from C1 to C3 were included in the reconstructed 3-D images. Gold color. GFP signal. Blue, autofluorescence. Boxed regions were enlarged in A1 and A2. (B). A sub-block of 500 µm thickness was displayed to show the dense projections of DRG neurons in the spinal cord. (C). Boxed regions in (B) were enlarged to show details. Boxed regions in (C) were enlarged or resliced in C1, C2, C3 and C4 to show axon arbor in C. Axonal bouton could be clearly visualized (arrows in C2 and C4). (D and E). Y-Z view and X-Z view of the image stack was displayed. Boxed regions in (E) were resliced or enlarged in E1, E2 and E3 respectively. Dotted lines in E2 and E3 indicated boundaries between two stitched adjacent stacks in z-dimension.

A human bone sample (1mX1mmX1mm) was stained with FITC to reveal the osteocytes and Haversian Canal system. Sample was processed following the TESOS method for hard tissue organs and imaged on a Zeiss upright two-photon microscope with 40X/1.3NA objective. The final image was stitched from 6 stacks in Z dimension (**Figure S3 A-C**). Optical slices acquired on various depth displayed equally high resolution (**Figure S3 D**). The Haversian Canal system with surrounding osteocytes were revealed through FITC staining (**Figure S3 E, F**). X-Z and Y-Z optical slices indicated continuous structure in Z dimension. Sectioning did not distort cellular organization (**Figure S3 H1**).

*Shh-Cre*^*ERT2*^*;Ai140* mice were used to label DRG neurons within spinal cord specifically. Adult mice were induced with tamoxifen and the cervical vertebrae segment including both spinal cord and surrounding bones were harvested. After transparent embedding, the sample was imaged with Leica 40X/1.3NA objective at 0.37 µm X 0.37 µm X 1.2 µm voxel size. We performed linear channel unmixing to distinguish autofluorescence from GFP signal. The final image stack (3.5 mm X 2.2 mm X 3 mm) was stitched from 22 stacks in Z dimension (**Figure 3 A, Video 2**). Fine neural structures were clearly visualized, including the DRG neuron soma (**Figure 3 A1**), central axonal branch (**Figure 3 A2**), axonal arbor (**Figure 3B, C, C1**) and boutons (**Figure 3 C1-C4**). The image was continuous on Z dimension (**Figure 3 D, E**). Fine structures including arbor and axons remains intact and continuous in Z dimension (**Figure 3E1-E3**).

These results indicated that TESOS method is applicable for organs of various tissue types and enabled sub-micron resolution imaging for large samples.

### TESOS method is compatible with immunofluorescence staining

We further tested if the TESOS method is compatible with immunofluorescent staining. Thick brain slice (1.5mm X 1.5mm X 1.0mm) of adult mice were performed wholemount immunofluorescent staining with antibodies against laminin for vasculature labelling. Samples were processed with the TESOS and imaged with a 40×1.3NA objective on a confocal microscope method in combination with sectioning microtome. Final image stack was stitched from 6 stacks in Z dimension (**Figure S4 A**). X-Y optical slices acquired at various depth showed comparable staining signal of vasculature at all depth (**Figure S4 B, B1, C, C2, D, D1, E, E1**). Sub-block on x-z dimension showed continuous vasculature staining signal in Z dimension (**Figure S4 F**). X-Z optical slice also showed that vasculatures staining signal was consistent and the structure remained continuous at all stitching boundaries (**Figure S4 G, G1-G6**). We also performed double immunofluorescent staining for mouse brain slice with antibodies against laminin and GFAP. The vasculature and glial cells were clearly displayed (**Figure S4 I, I1-I3**), which indicated that the TESOS method is compatible with immunofluorescent staining.

### Whole-body imaging of mouse pup at micron-scale resolution

*Thy1-YFP-16* mouse pup of P5 age was collected. The brain and internal organs were removed to facilitate penetration using an immersion protocol. All other tissues remained intact including skin, muscle, bones, eyeballs etc. (**Figure S1 C**). Samples were processed following TESOS method for hard tissue organs. The body trunk turned transparent after 2 weeks and remained equally transparent after transparent embedding initiated with UV light (**Figure S1 C**).

The sample was imaged with a Leica 20X/0.95 immersion objective on a Leica confocal microscope. The milling motor platform was used to remove surface tissue. The final image stack of 3.5cm(x) X1.0cm (y) X1.8cm (z) size was stitched from 40 stacks with ∼500µm thickness for each stack (**Figure 4A, F, Video 3**). The voxel size was 0.9 µm X 0.9 µm X 3.5 µm. Optical sections were acquired at various depth showing consistent resolution throughout the entire sample (**Figure 4 B, C**). At 1mm Z depth, neuron somas of the spiral ganglion were clearly visualized together which their axons innervating hair cells (**Figure 4 B, B1, B2**). In addition, retina, optic nerves, and trigeminal ganglions were also clearly displayed with soma and axon bundles being clearly shown (**Figure 4 B3, B4**). At 6mm Z depth, thoracic segment of the spinal cord was displayed. Enlarged images clearly showed the thoracic DRG neurons and the neuromuscular junction (NMJ) structures (**Figure 4 C, C1-C4**). To display the structure continuity in Z dimension, the image stack was displayed in lateral view (**Figure 4 D**) and Y-Z or X-Z optical slices were acquired at different locations. Y-Z optical slice showed 4 cervical DRGs (**Figure 4 E**). Y-Z or X-Z slices also displayed intact structures of L1 DRG, branchial plexus and sciatic nerves (**Figure 4 F, G, H**).

**Figure 4.**
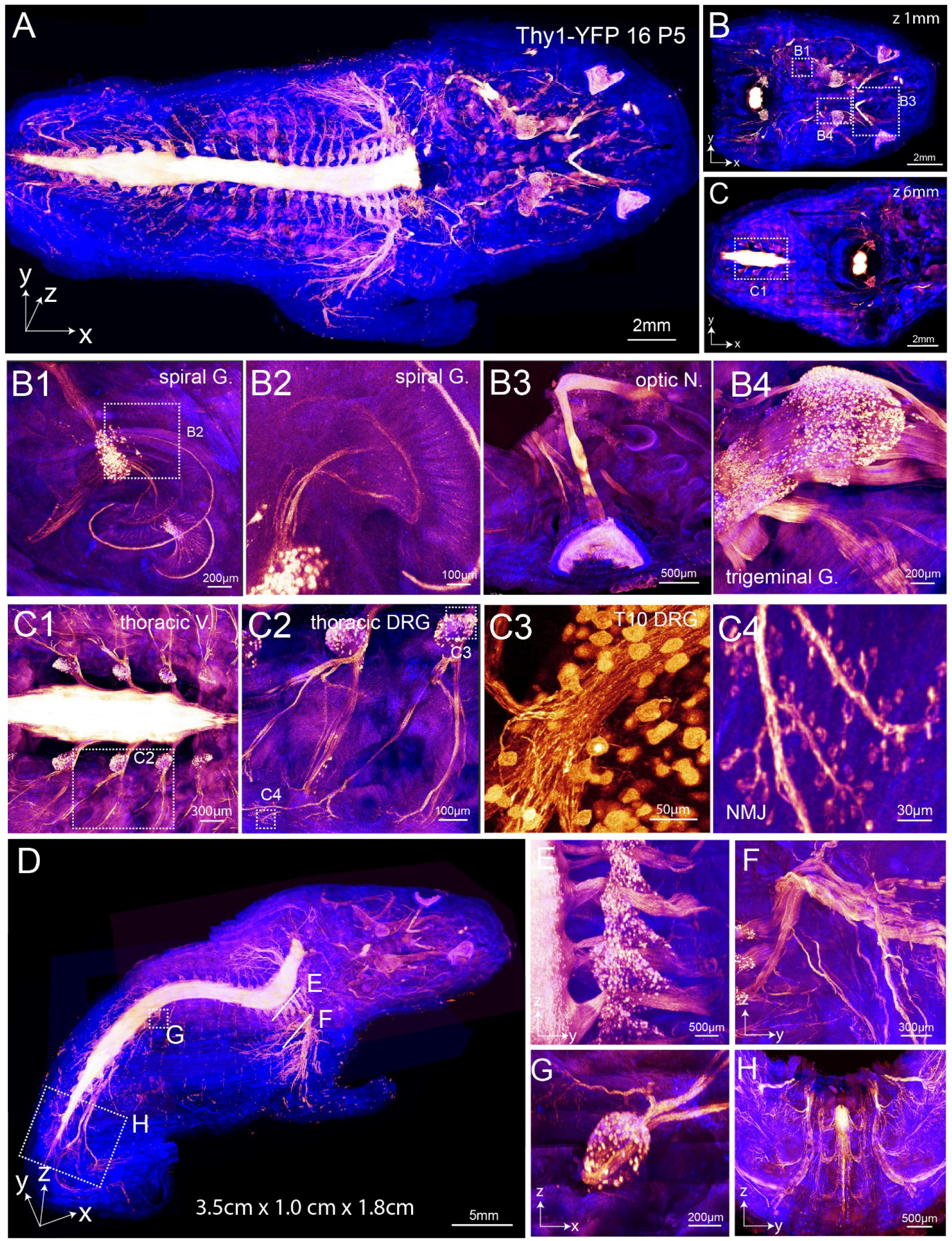
Whole body imaging at micron scale resolution. Brain and internal organs were removed from a *Thy1-YFP-16* mouse pup of P5. The body trunk was processed and imaged with a 20X 0.95NA oil immersion objective (gold, YFP signal; blue, autofluorescence). (A). Final image stack of 3.5cm(x) X1.0cm (y) X1.8cm (z) was stitched from 40 stacks with ∼0.5mm thickness for each stack. Voxel size is 0.9 µm X 0.9 µm X 3.5 µm. (B-C). Image in (A) was re-sliced at 1mm (B) or 6mm (C) Z-depth. Boxed regions were enlarged to display details. (B1). Spiral ganglion. Boxed region was enlarged in (B2) to show axons innervating hair cells. (B3). Optic nerve and retina. (B4). Trigeminal ganglion. (C1). Thoracic vertebrae. Boxed region was enlarged in (C2) to show two thoracis ganglions. Boxed regions in (C2) were enlarged to show the T10 DRG (C3) and the NMJ (C4). (D-H). Lateral view of the image stack (D). Boxed regions were resliced in Y-Z dimension to show the thoracis DRGs (E), branchial plexus (F), DRG T10 (G) and the sciatic nerve (H).

Therefore, the TESOS method in combination with milling platform enabled the micron-scale resolution imaging of whole body composed of various tissue types.

### Reconstruction of sensory field of individual sensory nerve axons on intact mouse paw

*Thy1-YFP16* mouse strain was known to label part of low threshold mechanical receptor (LTMR) neurons and endings of hairy skin and Meissner corpuscles of glabrous skin (Taylor-Clark et al., 2015). Intact forepaw of an adult *Thy1-YFP16* mouse was processed with the TESOS method for hard tissue organs. All tissue including hair, skin, muscle, bones and nerves was preserved. The sample was imaged with a Leica 40X/1.3NA objective on a Leica Sp8 confocal microscope. A sectioning microtome was used to remove surface tissue. The final image stack of 3.4mm(x)-4mm(y)-7.3mm(z) dimension was stitched from 35 stacks (**Figure 5A, Video 4**). The voxel size was 0.4 µm X0.4 µm X1.2 µm. Linear channel unmixing was used to distinguish GFP signal from autofluorescence derived from skin and skeletal muscles. A sub-block of 500µm z depth was displayed (**Figure 5 B**). Boxed regions were enlarged or re-sliced to display different types of nerve endings including Meissner corpuscles within the walking pad (**Figure 5 B1**), lanceolate ending surround the hair (**Figure 5 B2**), NMJ (**Figure 5 B3**), nerve axons in longitudinal direction (**Figure 5 B4**), cross section of nerve axons (**Figure 5 B5**) and branches of sensory nerve axon under the skin (**Figure 5 B6**). To display the tissue integrity and continuity, optical slice was acquired in Y-Z dimension (**Figure 5 C**). Boxed regions were enlarged to display sensory nerves innervating a walking pad (**Figure 5 C1**), NMJ on the muscle (**Figure 5 C2**) and LTMR nerve axons under the hairy skin (**Figure 5 C3, C4**). There are three carpal vibrissae located at the wrist of the forearm which sense the forearm position (**Figure 5A**) (Niederschuh et al., 2015). We were able to visualized nerve axons innervating the three carpal vibrissae (asterisks in **Figure 5 D**) (Niederschuh et al., 2015).

**Figure 5.**
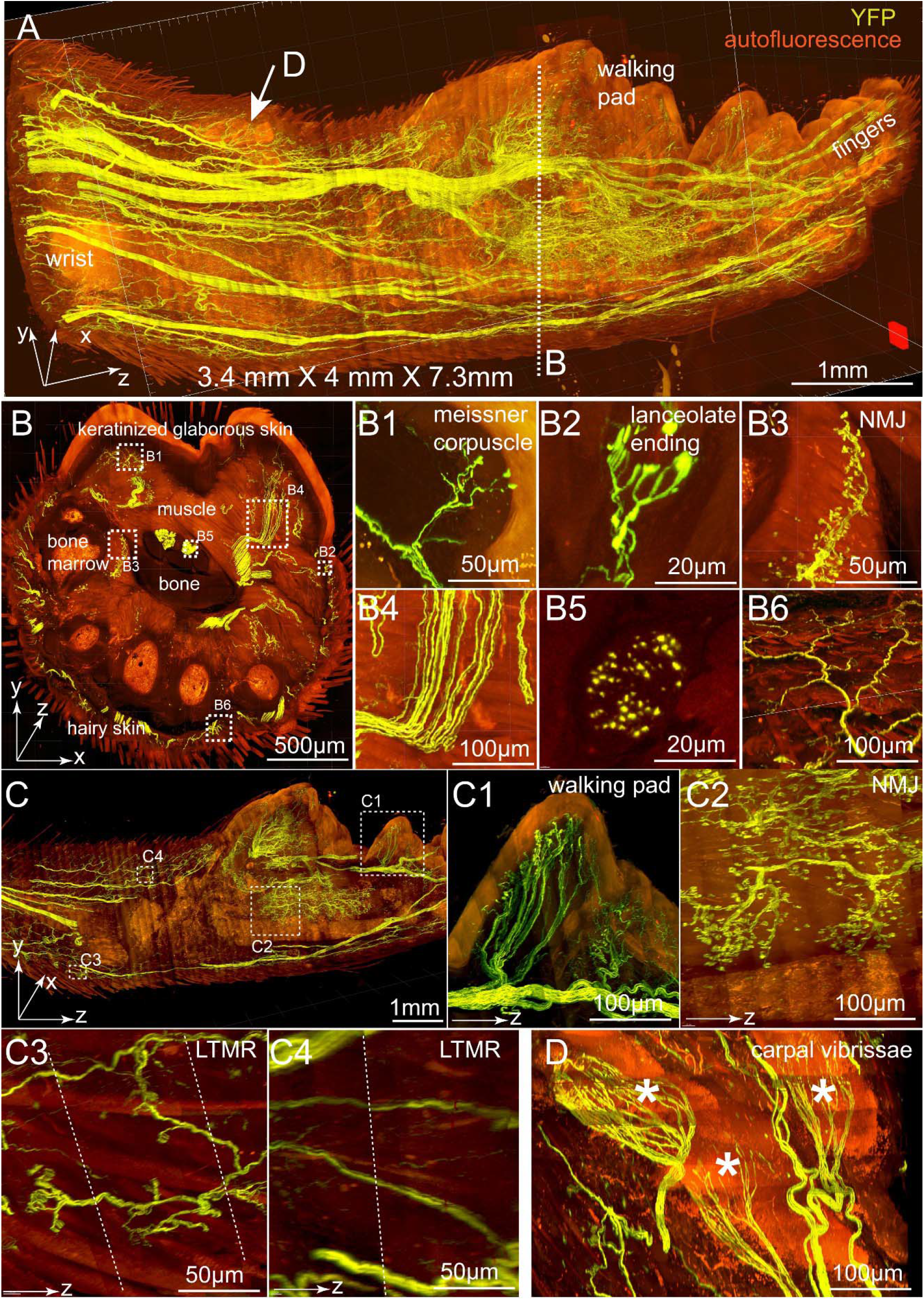
Nerves within the forepaw of an adult *Thy1-YFP16* mouse. An intact forepaw including the skin and hair was acquired from an adult *Thy1-YFP16* mouse. The sample was processed and imaged with a 40X/1.3NA objective (voxel size 0.4 µm X 0.4 µm X 1.2 µm). The digits were not included in the image. Yellow, YFP signal; Red, autofluorescence. (A). Reconstructed final image stack of 3.4mm(x)-4mm(y)-7.3mm(z) dimension was stitched from 35 stacks with 200-240µm thickness for each stack. (B). A sub-block of 500 µm thickness in z (dotted line in A), was displayed in x-y orientation. Boxed regions were enlarged or re-sliced to display various structures including Meissner corpuscle (B1), lanceolate ending surrounding a hair follicle (B2), neuromuscular junction (NMJ) (B3), nerve bundles in longitudinal dimension (B4), nerve bundles cross section (B5) and a sensory axon with its branches under the skin (B6). (C). A sub-block of 200 µm thickness in x was displayed in y-z orientation to display the structure continuity in the z-dimension. Boxed regions were enlarged or re-sliced to display various structures including sensory nerves within a walking pad (C1), NMJ (C2), LTMR innervating hair follicles (C3) and LTMR axons (C4). Dotted lines in (C3) and (C4) indicated boundaries between two stitched adjacent stacks in z-dimension. (D). Sensory nerves innervating the three carpal vibrissae (asterisks) at the wrist position (arrow in A). See also Figure S5.

The image stack was analyzed in Vaa3D. Nerve axons and endings were visualized within the glabrous skin of walking pad (**Figure S5 A**). All the axons under the glabrous skin of walking pad are sensory axons innervating Meissner corpuscles based on their morphology and localization within the dermal papillae (**Figure S5 A’**). All endings derived from individual axon could be traced (**Figure S5 B**). We traced all the labelled axons within the five walking pads on mouse forepaw (**Figure S5 C, D**). Around 5-15 sensory axons were labelled within each walking pad (**Figure S5 E1-E5**). Each axon innervated 7.2 ± 3.14 Meissner corpuscles on average. The average area of receptive field for each axon is 8620 ± 7820.80 µm^2^ (**Figure S5 F**). The receptive fields between axons did not overlap (**Figure S5 G**).

### Reconstruction and mapping of complete projection of single DRG neurons within the spinal cord

*Synapsin-Cre* AAV was injected under the glabrous skin of the walking pads of *Ai140* mice forepaws. Different AAV titers were provided for left or right side. Two months later, the complete cervical and thoracic vertebrae segments including spinal cord and surrounding tissues were collected to preserve intact DRG to spinal cord connections. Transparently embedded samples were imaged with a 40X/1.3NA objective under a Leica Sp8 confocal microscope. Microtome sectioning was used to remove surface tissue. Final image stack of 2.5mmX3.8mmX6mm was reconstructed from 28 stacks in z dimension. The voxel size was 0.4 µm X0.4 µm X1.2 µm.

All labelled neurons were located between DRGs C5-C8. Central projections were mostly between C4 and C8 segment (**Figure 6 A, Video 5**). X-Y sub-block showed the central projections are restricted to the medial side of the spinal cord (**Figure 6 B**). DRG neurons of various sizes were labelled with no obvious spatial pattern (**Figure 6 B1, B2**). All parts of the neuron were visualized, including DRG central axons, collateral branches, arbors and boutons (**Figure 6 B1, B2, C, C1, C2**). A motor neuron was also labelled with dendrites and axon clearly displayed (**Figure 6 D**). Lateral view of the image stack indicated that sectioning process did not disrupt the structural continuity. Reconstructed neuron soma and axon remained intact and continuous in z dimension (**Figure 6 E, E1, E2**).

**Figure 6.**
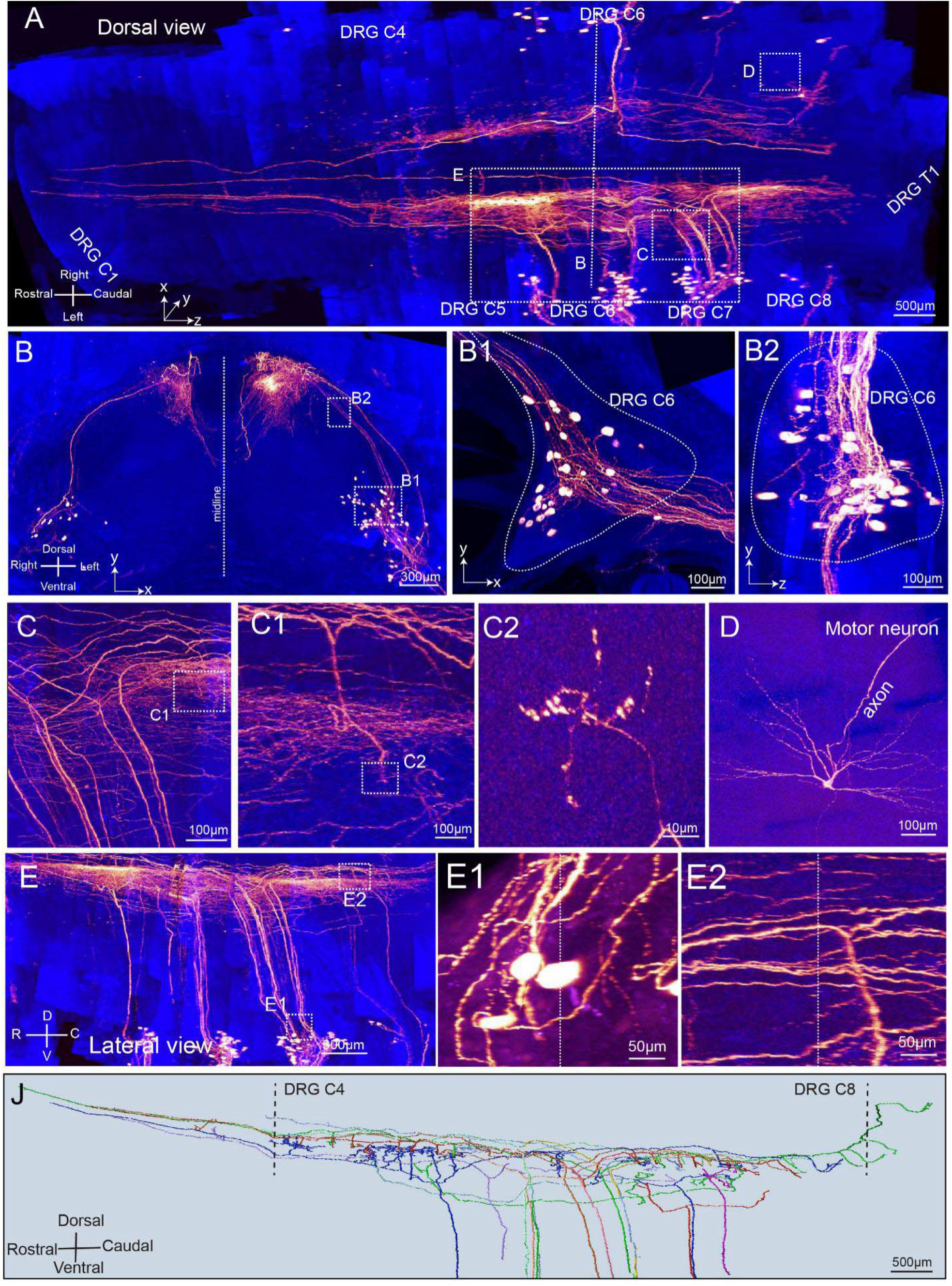
Complete projections of single sensory neurons in the spinal cord. DRG neurons were labelled by injecting adult *Ai140* mice with *Synapsin-Cre* AAV at the walking pad position. Lower AAV dosage was given to the right side than the left side. Cervical vertebrae was collected two months after injection and processed for imaging with a 40X/1.3NA objective (voxel size 0.4 µm X 0.4 µm X 1.2 µm). (Gold, GFP signal; Blue, autofluorescence). (A). Reconstructed final image stack of 2.5mmX3.8mmX6mm dimension was constructed from 28 stacks with 200-240µm thickness for each stack. (B). Caudal view of the image stack. Boxed regions were enlarged or re-sliced to display details including DRG neurons (B1, B2), (C). Bifurcations of DRG neuron central branches. Boxed regions in (C) were enlarged to display axon arbor (C1) and boutons (C2). (D). A motor neuron that was randomly labelled. (E). Lateral view of the image stack. Boxed regions were enlarged to display DRG neurons (E1) and axons (E2). Dotted lines in E1 and E2 indicated boundaries between two stitched adjacent stacks in z-dimension. (J). Complete projection of 12 DRG neurons within the spinal cord. See also Figures S6 and S7.

All fine details of single neurons could be identified (**Figure S6 A**). A sensory neuron in the DRG C6 gave rise to central and peripheral axons (**Figure S6 B**). The central axon bifurcated to form a caudal and a rostral branch (**Figure S6 C**). The caudal branch extended from DRG C6 to DRG C8, gave rise to 3 collateral branches with arbors and ended with an arbor (**Figure S6 D**). The rostral branch extended from DRG C6 to DRG C2, gave rise to 7 collateral branches with arbors and ended as a free terminal with no arbor (**Figure S6 D’**). All the 10 collateral branches with their arbors were displayed (**Figure S6 E1-E10**).

Sparse labeling and high-resolution imaging enabled us to outline and trace axon arbors in its entirety with Vaa3D (**Figure S7 A**). We were able to visualize the spatial overlap between central projection arbors of adjacent collateral branches from the same neuron (**Figure S7 B**). We also visualized spatial overlap between projection arbors from two different DRG neurons (**Figure S7 C**).

We outlined the complete central projection arbors of 12 DRG neurons (**Figure 6 J, Figure S7**). The findings are summarized in below: (1) Most of the arbors are localized between DRG C4 and T1 segment (10/12) (**Figure S7 D, E, F, G, H, I, K, L, N, O**). No arbor was visualized rostrally beyond DRG C2. (2). The axon arbor all projected into medial side of the spinal cord on laminae 3,4 or 5 (**Figure S7 D’-O’**). (3). The collateral branches and arbors are not evenly distributed along the axons. Rostral branches can extend 2-3mm without collateral branches or arbors (**Figure S7 D, E, G, I, K, L, M**) (4). None of rostral branches extended beyond DRG C1 position into the brainstem. Most of them (9/12) ended as free terminal without arbor (**Figure S7 D, E, G, H, I, J, K, L, M**). (5). Most caudal branches ended as arbors (9/12) (**Figure S7 D, E, F, G, H, I, J, K, M**).

### Reconstruction of long-range projection of sensory neurons from forepaw to spinal cord

Sensory neurons in the DRGs send out axons to both peripheral organs and spinal cord spanning a very long range. To outline the complete sensory neuron projections, adult *Thy1-EGFP* mice were used. The forepaw, forearm, cervical and thoracic vertebrae were isolated with all tissues in place. Samples were processed following the TESOS method for hard tissue organs. Processing time for each step was doubled to assure complete penetration. Samples were transparently embedded and imaged with a 40X/1.3NA objective on a Leica Sp8 confocal microscope with the voxel size 0.4 µm X 0.4 µm X 1.5 µm. The milling motor setup was used for removing surface tissue.

The final image stack (2.5 cm X 1.8 cm X 2 cm) was stitched from 98 stacks including the entire forepaw, the radial nerve, part of branchial plexus, DRG and spinal cord (**Figure 7 A, Video 6**). Comparing with *Thy1-EYFP-H* or *Thy1-YFP16* mouse strains, neuron labelling of *Thy1-EGFP* mice was sparser. LTMR sensory endings and axons were recognized through lanceolate endings surrounding hair follicles (**Figure 7 A**). The forearm section was selectively imaged surrounding the radial nerve. Small portion of nerve axons within the radial nerve was labelled which made the axon tracing possible (**Figure 7 A, D1, D2, D3, D4**). Spinal cord segment between C2 and T2 was imaged.

**Figure 7.**
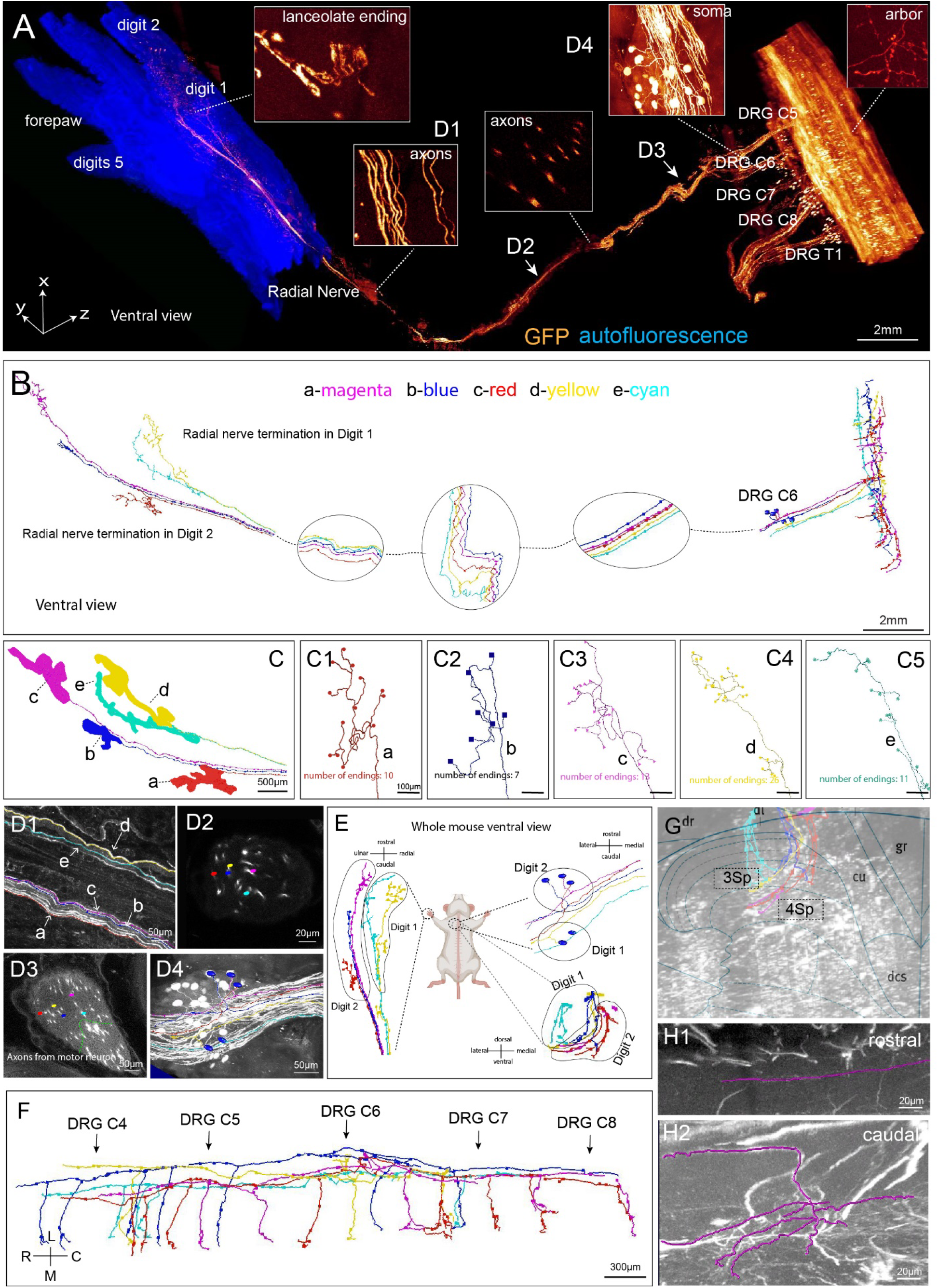
Mapping of individual sensory neurons from the hair follicles on the forepaw to the spinal cord. A body trunk spanning from the forepaw to the vertebrae of an adult *Thy1-EGFP* mouse was dissected and processed following the TESOS method for hard tissue organs. Embedded sample was processed with a milling motor setup and imaged under a confocal microscope with a 40X/1.3NA objective. (voxel size 0.4 µm X 0.4 µm X 1.5 µm). (**A**). Reconstructed final image stack with 25mmX18mmX20mm dimension were stitched from 98 stacks with 200-240µm for each stack. Indicated regions were enlarged in the boxes. (**B**). Complete tracing of five sensory neurons from hair follicles to the spinal cord with Vaa3D. Each color represents one neuron. (**C**). Receptive fields of the five sensory neurons innervating hair follicles on digits 1 and 2 (labelled as **a-e** with different colors). The terminals were shown as round dots in (**C1-C5**). (**D1-D4**). Optical slices acquired at different positions of the tracing course. Positions were indicated in panel (**A**). (**E**). A diagram showing the topological relationship between the sensory receptive fields on the digits, neuron somas in the DRG and axon projections in the spinal cord. (**F**). Complete tracing courses of the five neurons within the spinal cord. (**G**). Axonal projections were mapped to the Allen Spinal Cord Atlas at cervical 7 (C7) position. (**H1-H2**). The rostral (**H1**) and caudal endings (**H2**) of an axon.

Axons tracing was performed with Vaa3D. Finally, we were able to reconstruct complete projection courses of five DRG neurons with confidence. The entire tracing courses were ∼5.5 cm from forepaw digits 1 and 2 to the DRGs and extended ∼5 mm in the spinal cord (**Figure 7 B**).

The receptive fields of the five LTMR neurons were longitudinal along the digit axis (**Figure 7 C**). Numbers of lanceolate ending ranged from 7 to 26 (**Figure 7 C1-C5**). The target axons were traced within the radial nerve all the way to the branchial plexus and to the DRG C6 (**Figure 7 D1-D4**). The spatial relationship of the five axons were not constant within the radial nerve. The neuron somas innervating digit 1 were localized more caudally than those of digit 2 (**Figure 7 E**). The projection fields were mostly in the middle of lamina 3 (3Sp) and lamina 4 (4Sp). Projection fields of digit 2 were localized medial than those of digit 1 (**Figure 7 G, E**). In the rostral-caudal direction, projections of these LTMR neurons extended from DRG C3 to DRG C8 and each of them gave rise to 3∼8 collateral branches (**Figure 7 F**). None of their rostral branches extended beyond C3. Their rostral branch ended as free terminal and caudal branches ended as arbors (**Figure 7 H1, H2**).

## Discussion

Hydrogel was commonly used in many biomedical applications including tissue clearing techniques. Low concentration hydrogel polymerized from acrylamide and water protects proteins from harsh detergent treatment (Chung et al., 2013; Treweek et al., 2015; Yang et al., 2014). Organogel is a gel composed of an organic solvent phase dispersed within a cross-linked polymer and was used for drug delivery or cosmetic products (Skilling et al., 2014; Tan et al., 2020; Yuan et al., 2019). TESOS method is its first application in the microscopic imaging field. While working on our previous PEGASOS method (Jing et al., 2018), we noticed by coincident benzyl benzoate solution may form transparent gel in the presence of polyethylene glycol diacrylate (PEG-DA), while the samples within still remained transparent. Following this clue, we tested many combinations and identified BB-BED formula. In this formula, benzyl benzoate (BB) is the solvent disperse phase. Bisphenol-A ethoxylate diacrylate (BED) is a dental monomer crosslinker known for its biosafety and strength(Fleisch et al., 2010; Xie et al., 2017). Both BB and BED have high RI (BB, 1.56; BED 1.55). The high RI helps to render both soft and hard tissues transparent. Quadrol keeps the medium basic pH to protect GFP fluorescence (Jing et al., 2018; Schwarz et al., 2015). 2-Hydroxy-4’-(2-hydroxyethoxy)-2-methylpropiophenone UV initiator was used to initiate the radical polymerization reactions. Comparing with heat-initiation, UV-initiation provides much better control over the reaction heat, which helps protect GFP activity.

We define *transparent embedding* as the process of embedding samples without losing tissue transparency and endogenous fluorescence. In many formulas we tested, samples lost their transparency after polymerization despite of excellent transparency in the solution form. A formula composed of BB and PEG-DA also achieved transparent embedding. However, its organogel was too soft to provide mechanical support.

Block face imaging methods, including Mouselight and fMOST, were designed to achieve axon level isotropic resolution within rodent brains (Economo et al., 2016; Li et al., 2010; Tao et al., 2012; Zheng et al., 2013). However, none of them was applicable for organs made of complex tissue types, mostly due to insufficient embedding support. For block face imaging, top surface of the sample was removed and then the block surface was imaged. To stitch image blocks before and after processing together, it is important that samples remained no distortion and reposition without tilting.

Organogel composed of BB and poly-BED is nearly two folds stronger than a paraffin wax used for histological embedding. A tissue sample composed of both soft and hard tissues embedded within could therefore resist sectioning/milling process without distortion. We designed two processing strategies for reducing sample surface. Sectioning strategy was based on regular rotary microtome. Embedded samples were repeatedly transferred between microtome and microscope. Magnetic kinematic base is the key component to ensure precise re-position. This strategy could be easily achieved with commercially available parts without major modification to the microscope. Although we only tested Leica and Zeiss microscope in this study, sectioning strategy can be adopted to any upright confocal/two photon microscopes. Milling motor strategy was designed for processing large samples. For larger samples, minor positioning errors may accumulate during processing and made stitching challenging. The industrial linear guide repositioned samples with much higher precision. Although both strategies were currently operated manually, they can both be automized in the future.

Strong autofluorescence from skin, muscles and bone was another major challenge for imaging peripheral organs of complexed tissues (Monici, 2005). Immunostaining with nanobody was used to boost the signal over autofluorescence (Cai et al., 2019). In our study, attributed to the excellent preservation of endogenous fluorescence, we were able to utilize linear channel unmixing method to subtract autofluorescence from the authenticate GFP signal without boosting antibodies (Chorvat et al., 2005; McRae et al., 2019). This strategy was applicable for strong reporter strains including *Thy1-EGFP-M, Thy1-EYFP-H, Thy1-YFP16* and *Ai140. Ai14, Ai57* and *Ai65* were applicable for brain imaging, but not for peripheral organs due to their relative weak fluorescence intensity.

Organization of LTMRs under hairy and glabrous skins were previously studied using flat mount skin (Kuehn et al., 2019; Neubarth et al., 2020). TESOS method enables the sub-micron resolution imaging of an intact mouse paw and provided spatial organization information of sensory axons. Analysis results are similar with previously described (Neubarth et al., 2020).

Central organization and process of the information input from the sensory organs is a fundamental question for neuroscientist. In the direct DC pathway model, individual LTMR neurons convey information directly into brainstem DCN, where the information is relayed (Johnson, 2001; Mountcastle, 1957; Niu et al., 2013).More recently, an LTMR-RZ model was proposed. In this model, hairy skin LTMR synapse reside mainly within the spinal cord dorsal zone with few synapses in the DCN, indicating the spinal cord interneurons as the relay center for integrating somatosensory input (Abraira et al., 2017; Bai et al., 2015; Li et al., 2011).

It remained unknown if glabrous skin sensory neurons follow the same projection pattern. Our tracing results of 12 DRG neurons within the spinal cord provided direct evidence and suggests that sensory neurons for glabrous skin fit in the same model. All the DRG neuron somas labelled by AAV injected under the walking pad were localized in DRG C5, C6, C7 and C8. Their arbor projections are mostly between C4 and T1 segment. None of their axon or arbor extends beyond C1 into the brainstem.

*Syn-Cre* AAV injected in the walking pad has no selectivity for neurons types and may label both nociceptive and non-nociceptive neurons. For convenience, large neurons were chosen for our analysis. Although small DRG neurons were also visualized in the DRGs, few arbors were visualized beyond C4 in the rostral direction, suggesting that central projection of nociceptive neurons may also follow the same model. It appears that most sensation information from the forepaw walking pad was projected to a restricted spinal cord segment between C4 and T1, but not to the brainstem DCN.

Mesoscale brain connectome mapping is the frontier of neuroscience research, which provided critical information on organization principles of long-range neural connectivity (Peng et al., 2020; Winnubst et al., 2019). Mesoscale connectome mapping has never been achieved for the peripheral nervous system (PNS). Sensory neurons project their axons from peripheral organs to the spinal cord across various tissue types. Our study provided the first full course tracing of individual sensory neurons from digits to spinal cord in the entirety. Five sensory neurons innervating hairy skin LTMRs were reconstructed from the digits to the spinal cord spanning over 6cm. *Thy1-EGFP-M* strain was chosen because of its strong GFP fluorescence and relative sparse neuron labelling comparing with *Thy1-YFP-H* or *Thy1-YFP16*. Although tracing results from five neurons were insufficient for extrapolating significant conclusions, the TESOS method will provide a powerful tool for PNS connectome mapping at mesoscale in the future.

## Author contribution

H.Z. conceived the concept. H.Z. and W.G. designed and supervised the study. Y.Y., Y.M., S.Z and Y. W. performed most of the experiments with equal contribution. H.P., provided technical support on using Vaa3D. All authors discussed the results and commented on the manuscript text.

## Acknowledgements

We thank Drs. Lynn Opperman and Kathy Svoboda from the TAMU College of Dentistry for the research support. We thank Mrs. Meng Zhang, Ms. Evelyn Zhao and Ms. Hayley Zhao for the family support. This study was supported by the startup funding from the Chinese Institute for Brain Research to Hu Zhao and Woo-Ping Ge.

## Methods

### Animals

Mice were purchased from the Jackson Lab with genotypes including *C57BL/6* (JAX # 000664), *Thy1-EGFP* (JAX# 007788), *Thy1-YFP-16*(JAX #003709), *Ai14* (JAX# 007908), *Ai 140* (JAX# 030220) and *Shh-Cre*^*ERT2*^ (JAX# 005623). Human femur bone samples were kindly provided by Dr. Jian Q. Feng of the Texas A&M University. All animal experiments were approved by the Institutional Animal Care and Use Committee of the Chinese Institute for Brain Research.

For tamoxifen treatment, tamoxifen was dissolved in corn oil at 20 mg/ml. The solution was kept at -20°C and delivered via intraperitoneal injection or oral gavage for postnatal treatments.

### Preparation of TESOS solutions

#### Decalcification medium

Decalcification solution was composed of 20% (w/v) Ethylenediaminetetraacetic acid (EDTA) (Sigma-Aldrich E5134) in water. Sodium hydroxide (Sigma-Aldrich) was used for adjusting the pH to 8.0.

#### Decolorization medium

Quadrol medium (Sigma-Aldrich 122262) was diluted with H_2_O to a final concentration of 25 % (w/v). Due to the very vicious property of Quadrol, weighing is more convenient than measuring the volume. Quadrol medium can be warmed in a 45°C water bath to increase flowability.

#### Gradient tB delipidation medium

*tert*-Butanol (tB) (Sigma-Aldrich 360538) was diluted with water to prepare gradient delipidation solutions: 30% (v/v), 50% (v/v) and 70% (v/v). Quadrol (Sigma-Aldrich 122262) was added with 5% (w/v) final concentration to adjust the solution pH over 9.5.

#### tB-Q *dehydration medium*

Dehydrating medium was composed of 70% (v/v) *tert*-Butanol and 30% (w/v) Quadrol (Sigma-Aldrich 122262).

#### BB-BED clearing medium (Refractive Index R.I. 1.552)

Unless indicated in the manuscript, the BB-BED clearing medium was composed of 47%(v/v) benzyl benzoate (BB) (Sigma-Aldrich W213802), 48% (v/v) of bisphenol-A ethoxylate diacrylate Mn 468 (BED468) (Sigma-Aldrich 413550), 5% (v/v) Quadrol (Sigma-Aldrich 122262) and then supplemented with 2% w/v of 2-Hydroxy-4’-(2-hydroxyethoxy)-2-methylpropiophenone (Sigma Aldrich 410896) as the UV initiator. Bisphenol-A ethoxylate diacrylate Mn 512 (BED512) (Sigma-Aldrich 412090) could also be used to replace BED 468 at the same ratio with similar mechanical properties. The BB-BED medium was a colorless liquid with low viscosity and could be preserved at room temperature in the dark.

### Perfusion and tissue preparation

For trans-cardiac perfusion, mice were anesthetized with an intraperitoneal injection of a combination of xylazine and ketamine anesthetics (Xylazine 10-12.5 mg/kg; Ketamine, 80-100 mg/kg body weight). Ice-cold heparin PBS of 50-100 ml (10U/ml heparin sodium in 0.01M PBS) was injected transcardially to flush the blood. 50ml 4% PFA (4% paraformaldehyde in 0.01M PBS, pH 7.4) was then injected transcardially for fixation.

For the whole-body tissue clearing procedure, brain, tongue and all internal organs were removed. The eyeball and skin were preserved. For immersion of individual organs, the organs were immersed in 4% PFA at room temperature for 24 hours before clearing.

### Fluorescein isothiocyanate (FITC) staining for human bone samples

FITC staining was performed as previously described (Ren et al., 2014). The human specimen obtained from femur cortical bone was cut to cubic size with a diamond saw. The bone sample was fixed in 70% ethanol for two days at room temperature with a one-time change of 70% ethanol. The samples were then dehydrated in 95% ethanol for one day and 100% ethanol for one day. Next, the samples were immersed in 1% FITC (Sigma, cat. no. F7250) in 100% ethanol solution for 48hrs. After staining, samples were washed with PBS solution for 24 hrs and processed following TESOS method for hard tissues.

### Skin viral labeling

Skin viral injection protocol was modified based on previous publication(Bloom et al., 2019; Kuehn et al., 2019). *Ai140* (Jax 030220) reporter mice of P6 were used for viral injection. High titer (>10^12^GC/ml) AAV2/1-hSyn-Cre-WPREhGH virus (Addgene 105553-AAV1) was diluted 1:8 or 1:4 in 0.9% saline and 1µl was injected under the forepaw skin. The virus was a gift from James M. Wilson (Addgene plasmid # 105553; RRID:Addgene_105553). Mice were euthanized 2 months after injection.

### Whole-mount immunofluorescent staining

The whole-mount immunofluorescent staining was performed as previously described (Jing et al., 2018). After 4% PFA fixation, samples were decolorized with 25% Quadrol for 1 day and then washed with the PBS solution for 30 minutes. Samples were immersed in the blocking solution composed of 10% dimethyl sulfoxide (Sigma-Aldrich 276855), 1X casein buffer (Vector, SP-5020) and 0.5% IgePal630 (Sigma-Aldrich 18896) for blocking overnight. After blocking, samples were stained with the primary antibody diluted with the blocking solution for at least 72 hours at 4°C. Following that, tissues were washed with PBS at room temperature for 24hrs and then immersed in the secondary antibodies diluted with the blocking solution for another three days at 4°C. After secondary antibody staining, samples were washed with PBS for 6 hours and then moved to the delipidation and dehydration solutions following the TESOS clearing procedure.

Antibodies used for whole-mount staining included rabbit anti-GFAP antibody (1:100, Abcam ab7260), rabbit anti-Laminin antibody (dilution 1:50, Sigma-Aldrich L9393), goat anti-rabbit IgG Alexa Fluor 488 (dilution 1:200, Thermo Fisher A11034).

### Quantification of fluorescence intensity

Half brain slices of 0.5mm thickness from adult Thy1-EGFP mice were used for quantification of fluorescence intensity change. Fluorescent images of each time point were taken with a Zeiss stereo fluorescence microscope (Zeiss AxioZoom.V16) with the same zoom factor and exposure time. The “integrated intensity” values were measured using Image J (NIH). Average signal intensity of samples after fixation was defined as initial intensity and was normalized as 1.00. The ratio of “integrated intensity” value of other time points to initial intensity was calculated to evaluate the fluorescence intensity change.

### Quantification of Relative Young’s modulus

For quantification of Young’s modulus, a column with diameter of 3.5mm and height of 10mm was formed with 3% agarose, wax, and BB-BED solvent, respectively. Fixed brain and cleared brain were cut into the same shape for test. Young’s modulus was measured with tabletop Uniaxial Testing Machine (TestResources, USA), with compression load at the speed of 0.02mm/second. The average Young’s modulus of fixed brain was normalized as 1.00, and the ratio of Young’s modulus of other tissue or materials to that of fixed brain was calculated as the “relative Young’s modulus”.

### TESOS clearing procedure

For samples containing hard tissues, 4% PFA fixation was performed at room temperature for 24 hrs and then samples were decalcified in 20% EDTA (pH 7.0) at 37°C temperature on a shaker for 4 days. Samples were then washed with distilled water for at least 30 mins to remove excessive EDTA. Samples were next decolorized with the Quadrol decolorization solution for two days at 37°C on a shaker. Samples were placed in gradient tB delipidation solutions for 1-2 days and then tB-Q for 2 days for dehydration. Finally, samples were immersed in the BB-BED medium in a 37°C shaker for at least one day until transparency was achieved.

For soft tissues organs, decalcification treatment was skipped. After 24 hours fixation, samples were treated with Quadrol decolorization solution for 2 day at 37°C. Samples were next treated with gradient delipidation solutions in a 37°C shaker for 1 to 2 days, followed by TBQ dehydration treatment for 1 to 2 days. Finally, samples were placed in the BB-BED clearing medium in a 37°C shaker for at least 1 day until final transparency being achieved.

Before transparent embedding, cleared samples could be preserved in the clearing medium at room temperature in the dark for months.

The time schedule for clearing different types of tissue with immersion method is summarized in the following table.

**Table 1.** PEGASOS immersion method time schedule.

|  | Soft tissue organs | Hard tissue | Large body trunk |
| --- | --- | --- | --- |
| 20% EDTA | none | 4 days | 7 days |
| 25% quadrol | 2 days | 2 days | 2 days |
| 30% tert-butanol | 4 hours | 4 hours | 1 day |
| 50% tert-butanol | 6 hours | 6 hours | 1 day |
| 70% tert-butanol | 1 days | 1 days | 1 day |
| tB-Q dehydration | 2 days | 2 days | 2-3 days |
| BB-BED clearing | 1 day | 1 day | 2 days |
| Total time | 6-7 days | 10.5 days | 16-19 days |

### Transparent embedding of cleared samples through UV-initiated polymerization

The transparent embedding could be performed as early as 2 days after the sample transparency being achieved.

For small samples including mouse internal organs, regular plastic syringes were modified as the curing chamber. The adaptor and bottom of regular plastic syringes of 10ml or 25ml were removed. Samples were placed in the syringe barrel with 2-5 ml of BB-BED medium. Make sure the sample is away from the barre wall.

Syringe containing the samples was placed under a high-power UV curing light (Thorlabs CS20K2). The typical power readout for curing a cleared mouse brain is 50mW/cm^2^, the lamp head was 8 cm from the samples. Curing time was 5 minutes. Such set up could achieve full polymerization without causing sample overheating (below 55°C), which could quench endogenous fluorescence signals.

Alternatively, inexpensive industrial UV light source purchased from eBay could also be used. The power setup and curing time need to be optimized to achieve polymerization without overheating. At any time, the temperature should not be over 55°C to preserve endogenous fluorescence.

After UV curing, the samples were removed from the syringe by pushing the plunger. A scalpel was used to trim extra gel. Make sure to preserve at least 3-5mm gel surrounding the sample to provide sufficient support during cutting. Embedded samples could be preserved in a 50ml tubes. Polymerized organogel has slightly higher RI (1.560) than the solvent solution (1.552).

For transparent embedding of much larger samples including mouse pup whole body or adult mouse body trunks, customized glass chambers were used as the curing chamber. The UV curing was performed from all angles and the curing time was determined based on preliminary experiments. Typically, a mouse pup whole body could be polymerized within 30 minutes. After removing embedded samples from the chamber, the samples could be furtherly illuminated with the UV light from every angle to ensure complete polymerization. A complete polymerization could be indicated when fingers holding the gel felt no more heat being generated.

Polymerized samples remained transparent as in the solvent solution. The sample transparency could reduce significantly 10 days after curing, possibly due to ongoing polymerization within the tissue. Therefore, embedded samples are better preserved at 4 °C in the dark and imaged as soon as possible.

### Mounting and alignment of embedded samples on a rotary microtome

Magnetic kinematic base (SB1/M, Thorlabs) was used for mounting samples. The kinematic base was designed for re-mounting optical components with high repeatability and high precision. The SB1(/M) round kinematic base is composed of a top plate and a bottom plate. The top plate can be removed and replaced with an ON/OFF switch bar which interrupts the magnetic force coupling the plates of the base.

Embedded samples were glued onto the top plate of the kinematic base (SB1/M, Thorlabs) with two-component 5-min cure epoxy glue (Gorilla, Home Depot). The bottom plate of the kinematic base was screwed onto a mounting base (Thorlabs BA1/M). The mounting base was tightly clamped onto the specimen clamp of any lab microtome (either Sakura ACCU-CUT SRM or Leica 2050 was used in our lab).

Another bottom plate was secured with a screw onto a 2-axis goniometer stage (Thorlabs GNL20/M). The goniometer stage was screwed onto the sample movement stage (Scientifica MMBP motorized XY stage in our lab) of the microscope (Leica Sp8 confocal). For other microscopy brands, we improvised customized design to connect the kinematic base bottom plate onto the sample stage.

The sample will be transferred between the kinematic base on the microtome or on the microscope stage. Alignment was performed prior to imaging a sample. An empty kinematic base top plate was locked onto the microtome bottom plate. The sectioning movement plane is adjusted by adjusting the block adjustment handles until it is parallel to the top plate surface.

After the microtome alignment, an embedded test sample was locked onto the microtome. Sectioning (5µm/cut) was performed to expose a large area of sample surface. The sectioned sample was transferred onto the kinematic base on the microscope stage. The samples were examined with a 10X objective. Endogenous fluorescence is not required for this step since the tissue can be visualized based on autofluorescence. A large tile scan was performed by moving the X-Y stage. The goniometer stage was adjusted until the sample sectioning plane is parallel to the XY stage moving plane.

The alignment needs to be performed only once for a sample and should never be changed till the entire imaging process is completed.

### Imaging and sectioning of the transparently embedded samples

Embedded samples were mounted on the microtome and sectioned (5 µm/cut) to expose the sample top surface. After sectioning, the surface was dropped with BB-BED medium and covered with a glass coverslip. Samples were next cured with a UV lamp (Thorlabs CS20K2) for five seconds to polymerize the newly added medium and to secure the coverslip.

The embedded samples could be imaged with an upright confocal/2-photon microscopy as a regular slide. For immersion objectives, immersion oil could be dropped directly onto the coverslip. Oil immersion objectives are highly recommended due to the high R.I. (1.55-1.56) of the BB-BED medium and gel. The imaging depth of each imaging Z-stack is determined by the objectives and imaging requirements. The top plane of the Z stack should be at least 10µm below the sample sectioning surface to avoid any potential distortion on the machined surface.

After imaging the selected sample areas, the coverslip was removed by sliding it off the surface. The sample was transferred to the microtome for sectioning (5µm/cut). The sectioning depth should be at least 10% less than the Z-stack depth to provide overlapping area for stacks stitching. Sectioned sample was repositioned onto the kinematic base on the microscopy stage and dropped with BB-BED medium followed by coverslip placement and UV curing for the next imaging cycle.

Such imaging-sectioning cycles were repeated until the region of interest (ROI) was completely imaged.

### Milling of transparently embedded samples with a milling motor platform

Very large samples including mouse pup whole body and adult mouse body trunk were processed with a milling motor setup built on an upright Leica Sp8 confocal microscope. A high precision linear guide (IKO LWLF 42B) was installed on the sample XY movement stage (Scientifica Inc. MMBP). One end of the linear guide was under the objective and the other end was under the milling platform. Under the objective, a kinematic magnetic base plate (SB1/M, Thorlabs) was connected with the sliding block (IKO LWLFF 42BCS) on the linear guide through a vertical translational stage (Thorlabs MVS 005). The vertical stage was equipped with a fine-adjustment micrometer (Thorlabs 148-811ST). The milling platform was installed on the left side of the confocal microscope and was composed of a water-cooled spindle motor (Huanyang 300W 60K RPM) installed on a vertical translational stage (Thorlabs VAP 10/M).

Transparent embedded samples were glued on to the kinematic base top plate and locked onto the base plate under the objective. The samples were first moved along the linear guide to under the milling motor. Sample surface was milled off with a 2mm end-cutting milling bur (eBay). The sample movement under the motor was controlled by the sample stage. The cutting depth was controlled by adjusting the vertical stage position with the micrometer. The debris was blown off and a drop of BB-BED medium was placed onto the milling surface and a coverslip was placed. After curing the newly added medium with a UV light, the sample was moved back to the objective along the linear guide. Mechanical stops were installed on the linear guide to ensure repeatable reposition. The top plane of the Z stack should be at least 25 µm below the milling surface due to the relatively higher roughness from the milling process.

No alignment was needed for the milling setup since the milling X-Y movement during the milling was controlled by the sample stage so that the milling surface was already in line with the imaging plane.

### Linear channel unmixing for autofluorescence reduction

For tissues with strong autofluorescence including skin, skeletal muscles and bones etc. linear channel unmixing was performed to distinguish true fluorescent signal from tissue autofluorescence. Autofluorescence signal widely spreads from 350nm to 600nm (Zipfel et al., 2003), whereas fluorescence from protein or antibody conjugation is very restricted. Depending on the endogenous fluorescence spectrum, the autofluorescence detection channel was setup at either 488nm or 568nm wavelength. The imaging parameters for autofluorescence detection channel were carefully adjusted so that little true fluorescence signal was detected.

The channel unmixing operation was performed with the “image calculator” function in the Image J (NIH Image J). The autofluorescence signal was subtracted from the true signal channel to generate a new channel and the new channel was combined with the previous autofluorescence channel to generate the final images. Autofluorescence signal was used for outlining tissues.

### Image deconvolution and stitching

Image deconvolution was accomplished with the “microvolution” module within the Slidebook 6.0 software (3I inc.). The iteration number was set as 10. Regularization was set as “none”. Blind deconvolution was checked. PSF model was automatically generated based on the optical parameter being provided.

BigStitcher module within the Image J was used for stitching imaging tiles. 10% overlapping was set up for adjacent tiles. Image stacks were exported as image sequence of .tif format. Image stacks before and after samples sectioning/milling were sequentially stitched based on the overlapping regions. For two consecutive stacks, to facilitate speed, only images sequence in the overlapping zone were selected. Pairwise stitching module of the Image J was run to stitch these selected images together. The other image sequences in the two stacks were modified their metadata (custom Image J macro code).

### Image reconstruction and axon tracing

For 3-D rendering, image sequences were converted into. IMS format (Imaris Converter, Bitplane). 3-D rendering, snapshots and animation were performed with Imaris (Bitplane). Multiresolution pyramid was generated from stitched image sequences for 3-D tracing (TeraConverter, Bria et al. 2016). Manual tracing of axons was performed by a team of two annotators using TeraFly in Vaa3D (Peng et al., 2014; Peng et al., 2010). For registration, spinal cord image stacks were resliced for planes orthogonal to the rostrocaudal axis of the spinal cord. The resliced imaging plane was aligned and annotated based on the spatial map for the Allen Mouse Spinal Cord Atlases (the Allen Institute).

### Axon tracing from the paw to the spinal cord

To reduce the final data size for axon tracing, the stitched imaging data was separated into three parts: 1. nerve ending in paw; 2. nerve bundle in the arm; 3. DRG and projection within the spinal cord. An overlap of 40 slices was retained in each part for the determination of axon positions. For the first part, manual 3D tracing of axons was performed with TeraFly in Vaa3d. For the second part, tracing of single axon was performed manually using stitched 2D slices in Image J. Target axon from the first part was shown in “Section” mode in Vaa3D in order to display its localization in 2D slices, and to find its counterpart in the second part. Finally, axon tracing into the DRG and its projection within the spinal cord was done manually with Terafly in Vaa3D. Each tracing procedure was performed by a team of two annotators.

### Quantification and statistical analysis

*N* number are reported in figures and legends. Data are presented as mean ± standard deviation using one-way ANOVA or Student’s *t* tests. Statistical analysis was performed in GraphPad Prism and Microsoft Excel.

## Notes

https://www.youtube.com/channel/UCtDCiJ_yAH8xAsTtK4CqTGw

### Competing Interest Statement

The authors have declared no competing interest.

