## supplementary data for "Mapping of individual sensory nerve axons from digits to spinal cord with the Transparent Embedding Solvent System"

High resolution supplementary videos (Videos 1-6) are also available online in the Youtube channel.

<https://www.youtube.com/channel/UCtDCiJ_yAH8xAsTtK4CqTGw>

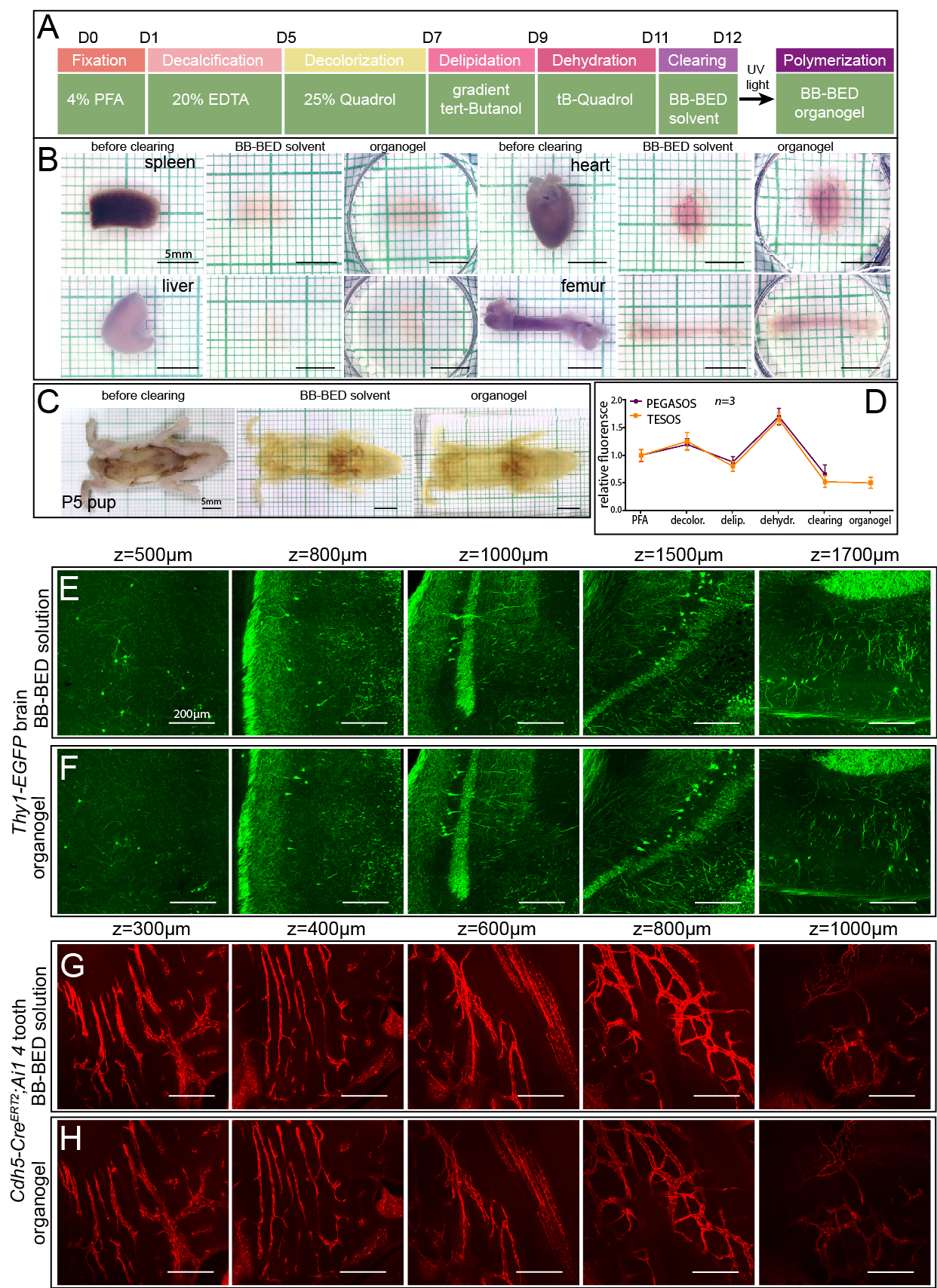
Figure S1.

**Figure S1. TESOS method enables transparent embedding of various tissue and organs without losing transparency and endogenous fluorescence**. **Related to Figure 1**.

(A). Treatment steps of TESOS method for mouse samples containing hard tissue.

(B). Images of spleen, heart, liver and femur harvested from adult mice before clearing, after clearing and after polymerization.

(C). A mouse pup of P5 with the brain and internal organs being removed was processed following the procedure listed in (A) and imaged before clearing, after clearing and after polymerization.

(D). Relative GFP fluorescence intensity at each step of TESOS treatment in comparison with the PEGASOS tissue clearing method.

(E, F). Thy1-EGFP mouse brain was processed with the TESOS method for soft tissue organs. Images were acquired with a 20X/0.95NA objective at various depth before (E) and after (F) organogel polymerization.

(G, H). Femur of adult *Cdh5-Cre^ERT2^;Ai14* mouse was processed with TESOS method for hard tissue organs. Images were acquired with a 20X/0.95NA objective at various depth before (G) and after (H) organogel polymerization.

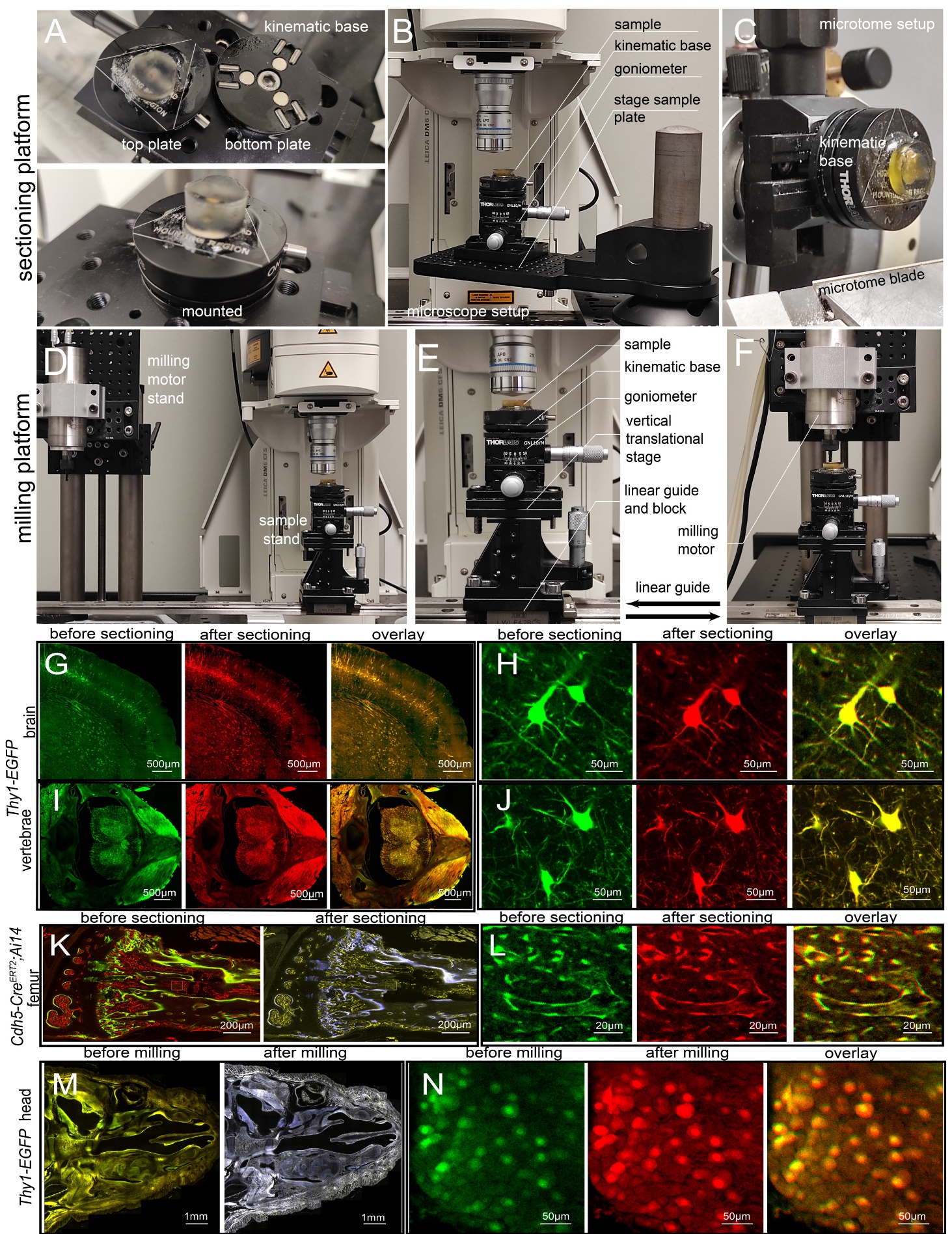
Figure S2.

**Figure S2. Microtome and milling platforms for sectioning or milling embedded samples.** **Related to Figure 1.** Kinematic magnetic base was used for sample mounting and transferring. Embedded sample was glued onto the top-plate of a kinematic magnetic base. Two bottom-plates were secured under microscope and microtome respectively. Top-plate with the sample was transferred between these two bottom plates. The magnetic design enables the top plate to automatically reposition to an exact location with high repeatability.

(A). Kinematic magnetic base with the two parts separated (top panel) or mounted (lower panel). (B). Description of parts when the sample was mounted under the microscope.

(C). Description of parts when the sample was transferred to the microtome.

On a milling platform, milling motor was built next to the microscope. The sample was moved between the microscope and milling motor along a high precision linear guide.

(D). Description of parts for the milling setup.

(E). Description of parts when the sample was under the microscope.

(F). Description of parts when the sample was under the milling motor.

(G-N). Transparent embedding enables sectioning or milling process with no detectable tissue distortion.

The brain of adult *Thy1-EGFP* mouse was processed and embedded following the TESOS method for soft tissue organs. The vertebrae, femur and the head samples were all harvested from adult mice and embedded following the TESOS method for hard tissue organs. Images were acquired with a 10X/0.4NA air or 20X/0.95NA immersion objective before and after tissue was sectioned/milled off. Imaging planes before sectioning/milling were 400µm below the surface. Samples of 390µm thickness were sectioned. Samples of 380µm thickness were milled off.

(G). The embedded brain was imaged before and after sectioning with 10X objective. Two images were overlaid to display unchanged morphology.

(H). Selected neurons were re-imaged with 20X objective.

(I). The vertebrae sample was imaged with 10X objective before and after sectioning and then overlaid.

(J). Selected neurons within spinal cord were imaged with 20X objective to display unchanged neurons morphology.

(K). Images of femur were acquired with 10X objective before and after sectioning.

(L). Selected regions in femur were re-imaged with 20X before/after sectioning.

(M). The head sample was imaged before and after milling.

(N). Selected ganglion neurons were re-imaged with 20X objective to display consistent morphology.

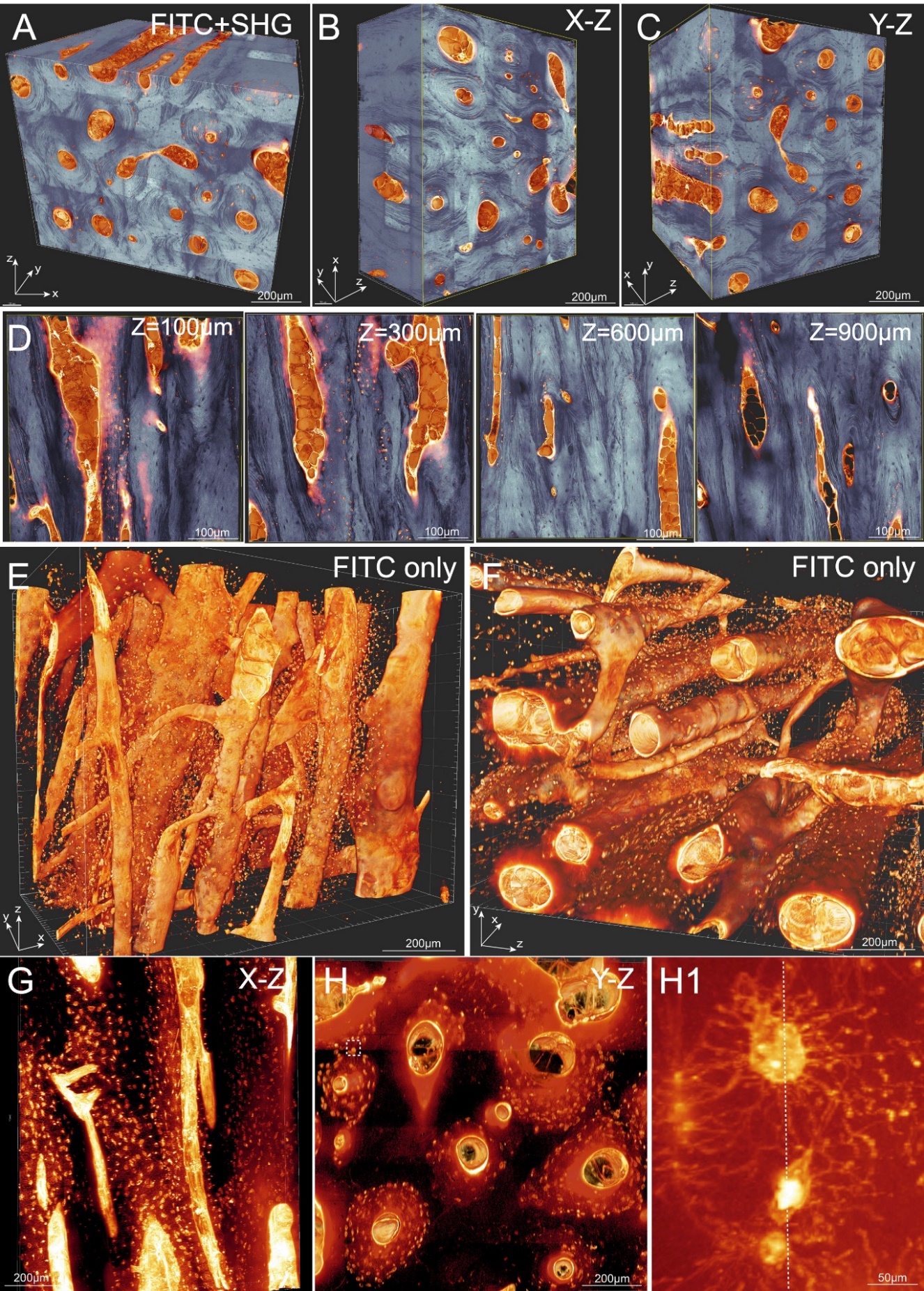

Figure S3.

**Figure S3. High resolution imaging of the human bone sample stained with fluorescein isothiocyanate dye (FITC). Related to Figure 2.** Human femur bone sample of 1mm (x) X 1mm (y) X 1mm (z) size was stained with FITC dye and processed following the TESOS method for hard tissue organs. Embedded bone was imaged under a Zeiss two-photon microscope with a 40X 1.3NA objective with the voxel size 0.4µm X 0.4µm X 1.2 µm. (Gold, FITC stain; light blue, second harmonic generation (SHG) signal).

(A) The final image stack was stitched from 6 stacks with 240µm thickness for each stack.

(B, C). Sub-blocks acquired in x-z (B) or y-z (C) dimension.

(D). Slices were acquired at various z-depth.

(E, F). FITC staining was displayed alone to show the Haversian Canals system in 3-dimension.

(G, H). x-z or y-z optical slices were acquired. Boxed region was resliced and enlarged in (H1). Dotted line indicated boundaries between two stitched adjacent stacks in z-dimension.

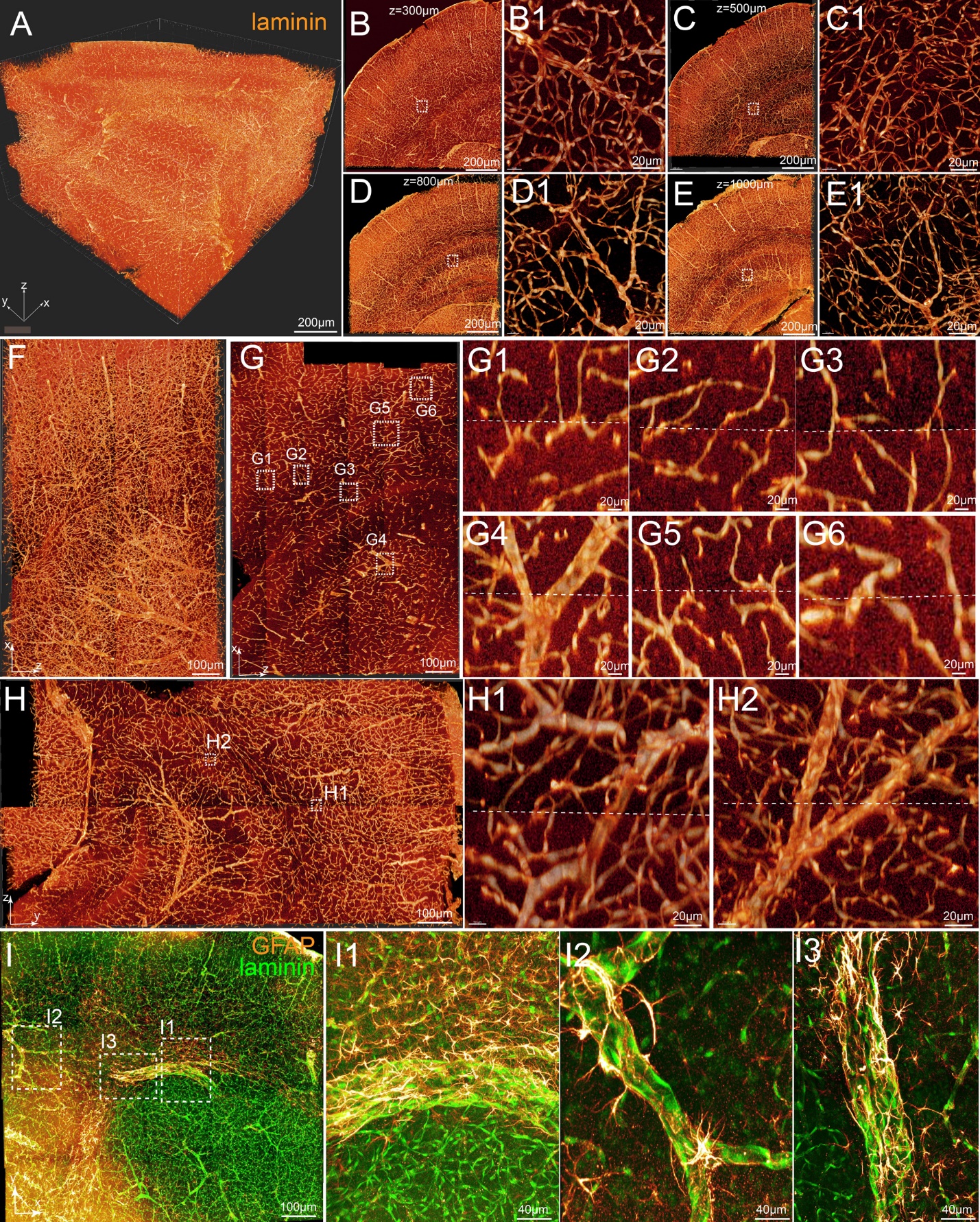

Figure S4.

**Figure S4. High resolution imaging of samples stained with antibodies.** **Related to Figure 2.** A piece of mouse brain with 1.5mmX1.5mmX1.0mm dimension was performed whole mount immunofluorescent staining with antibodies against laminin (A-H) or laminin + GFAP (I). Samples were processed with the TESOS method and imaged under a confocal microscope with a 40X/1.3NA objective. Voxel size is 0.4µm X 0.4µm X 1.2 µm.

(A). Reconstructed final image stack of 1.5mm(x)-1.5mm(y)-1.0mm(z) dimension was stitched from 6 stacks with 200-240µm thickness for each stack.

(B-E). Optical slices in x-y orientation were acquired at various depth. Boxed regions were enlarged (B1-E1).

(F) A sub-block of 1.5mm(x)-0.5mm(y)-1.0mm(z) was displayed in x-z orientation.

(G). An optical slice in x-z orientation. Boxed regions were selected at the boundary between adjacent z-stacks and enlarged in (G1-G6). Dotted lines indicated boundaries between two stitched adjacent stacks in z-dimension.

(H). An optical slice in y-z orientation. Boxed regions were enlarged in (H1) and (H2). Dotted lines indicated boundaries between two stitched adjacent stacks in z-dimension.

(I) Reconstructed final image stack for brain sample stained with GFAP + laminin antibodies. Boxed regions were enlarged in (I1-I3).

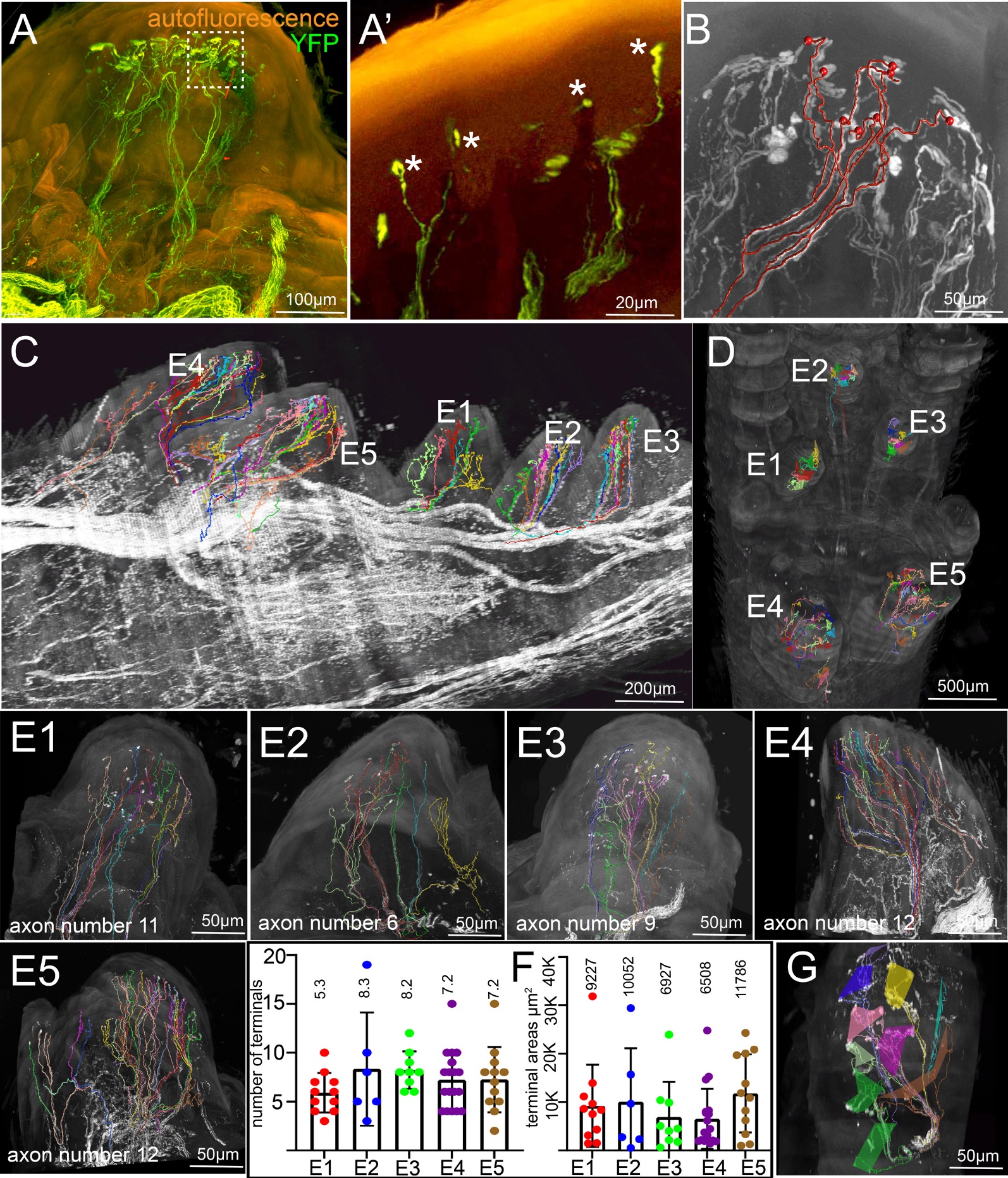
 Figure S5.

**Figure S5. Tracing of sensory axons innervating Meissner Corpuscles under the five walking pads of adult *Thy1-YFP16* mouse forepaw**. **Related to Figure 5**.

(**A**). A sub-block from the same image stack in **Figure 5** was acquired to display a walking pad and their innervating nerve axons. Boxed region was resliced in (**A’**) to show the morphology of the Meissner corpuscles (asterisks) and their localization within the dermal papillae.

(**B**). Tracing of one sensory axon and all of its derivative branches and terminals with Vaa3D.

(**C, D**). Tracing of all the sensory axons within the five walking pads (labelled as E1- E5) displayed from lateral (**C**) or ventral (**D**) view.

(**E1-E5**). Tracing results of the five walking pads was individually displayed.

(**F**). Quantification of average terminal number from one sensory axon (left panel) and average receptive field area of one sensory axon (right panel).

(**G**). The non-overlapping tiling pattern of different sensory axons receptive fields in walking pad E1.

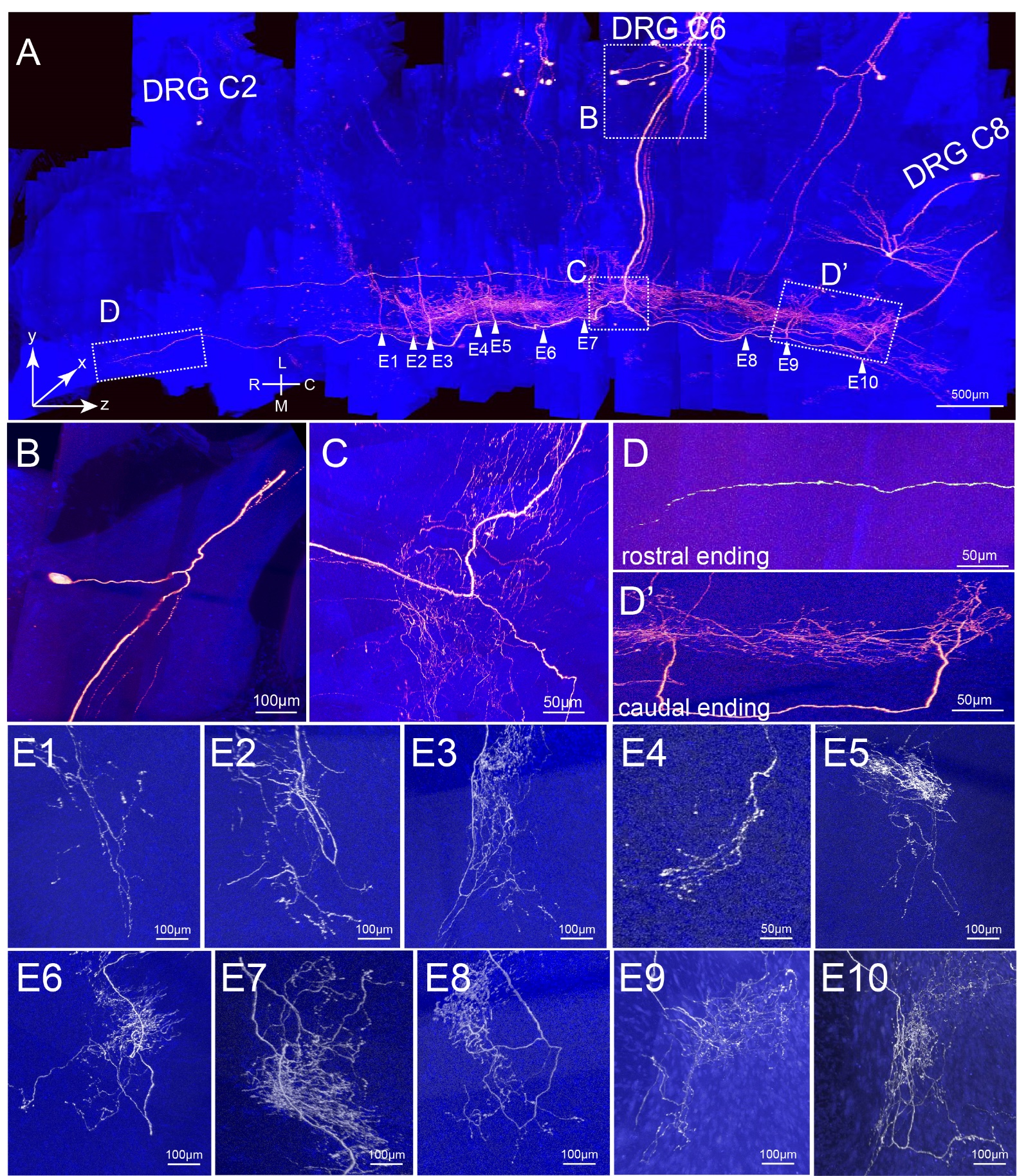
 Figure S6.

**Figure S6. Complete projection of one sensory neuron within the spinal cord.** **Related to Figure 6**. The sample was the same as the one in the Figure 6.

(**A**). A sensory neuron within C6 DRG and its projection within the spinal cord was displayed. Boxed regions were enlarged in following panels.

(**B**). The soma and the bifurcations of peripheral branch and central branch.

(**C**). The central axon branch bifurcated into caudal and rostral branches.

(**D, D’**). The rostral (**D**) and caudal (**D’**) termination of the axon.

(**E1-E10**). The 10 collateral branches and their arbors. The locations were indicated in (**A**).

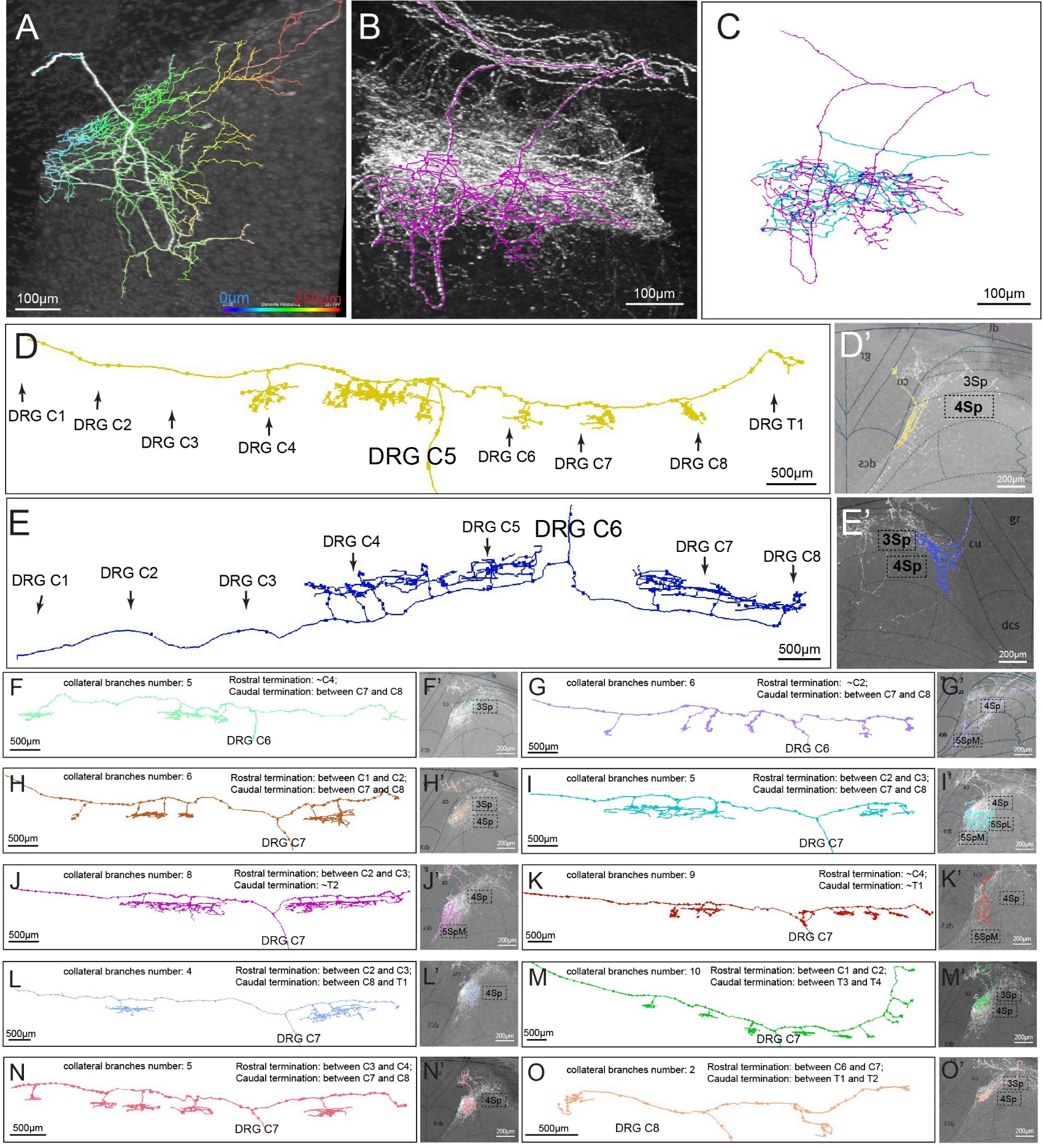
 Figure S7.

**Figure S7. Complete projection mapping of twelve sensory neurons within the spinal cord**. **Related to Figure 6**. Axon arbors were traced with Vaa3D.

(**A**). Color coded depth showing the spatial distribution pattern of an axon arbor.

(**B**). Arbors from two adjacent collateral branches derived from the same neuron.

(**C**). Spatial overlapping of arbors from two neurons.

(**D-O**). Complete tracing of 12 sensory neurons within the spinal cord. Eleven were from the left DRGs C5-C8 (**E-O**) and one was from the right side DRG C6 (**D**).

(**D’-O’**). Projections of arbors were mapped with the Allen spinal cord atlas. Boxed label indicated projection laminae. Collateral branches number, positions of rostral and caudal terminations were described in each figure.
